# Decreased miR-24-3p potentiates DNA damage responses and increases susceptibility to COPD

**DOI:** 10.1101/2020.05.22.108688

**Authors:** Jessica Nouws, Feng Wan, Eric Finnemore, Willy Roque, Sojin Kim, Isabel Bazan, Chuan-xing Li, C. Magnus Skold, Xiting Yan, Veronique Neumeister, Clemente J. Britto, Joann Sweasy, Ranjit Bindra, Åsa M. Wheelock, Jose Gomez-Villalobos, Naftali Kaminski, Patty J. Lee, Maor Sauler

**Affiliations:** Section of Pulmonary, Critical Care, and Sleep Medicine, Department of Internal Medicine, Yale University School of Medicine, New Haven, CT, USA; Department of Anatomy, Beijing University of Chinese Medicine, Beijing, China; Department of Internal Medicine, Rutgers New Jersey Medical School, Newark, NJ, USA; Division of Respiratory Medicine and Allergy, Department of Medicine and Center for Molecular Medicine, Karolinska Institutet and Karolinska University Hospital, Stockholm, Sweden; Department of Pathology, Yale University School of Medicine, New Haven, CT, USA; Department of Radiation Oncology, University of Arizona College of Medicine, Tucson, AZ, USA; Department of Therapeutic Radiology, Yale University School of Medicine, New Haven, CT, USA; Section of Pulmonary Allergy and Critical Care Medicine, Department of Internal Medicine, Duke University School of Medicine, Durham, NC, USA

## Abstract

Activation of the DNA damage response (DDR) due to chronic exposure to cigarette smoke (CS) is implicated in the pathogenesis of Chronic Obstructive Pulmonary Disease (COPD). However, not all smokers develop COPD and the pathologic consequences of CS exposure are heterogenous. Cellular mechanisms that regulate the DDR and contribute to disease progression in susceptible individuals are poorly understood. Because microRNAs are well known regulators of the DDR, we evaluated microRNA expression arrays performed on lung samples from 172 subjects with and without COPD. We identified miR-24-3p as the microRNA best correlated with radiographic emphysema (ρ=-0.353, P=1.3e-04) and validated this finding in multiple cohorts. In a CS-exposure mouse model, miR-24-3p inhibition increased emphysema severity. In human airway epithelial cells, miR-24-3p suppressed apoptosis through the BH3-only protein BIM and suppressed homology-directed DNA repair and the DNA repair protein BRCA1. Finally, we found BIM and BRCA1 were increased in COPD lung tissue and inversely correlated with miR-24-3p expression. We concluded that decreased miR-24-3p expression increases COPD susceptibility and potentiates the DDR through BIM and BRCA1.

## INTRODUCTION

Chronic Obstructive Pulmonary Disease (COPD) is a leading cause of global mortality, characterized by persistent airflow limitation due to small airway disease and emphysema (1). Chronic exposure to cigarette smoke (CS) is a major risk factor for COPD and cytotoxic effects of CS contribute to COPD pathogenesis. However, only certain smokers develop clinically significant COPD. Therefore, individual differences in cellular stress responses to CS may be critical determinants of COPD pathogenesis (2, 3). Currently, there is limited understanding of the specific cellular stress responses that predispose or protect individuals from disease progression in COPD.

DNA damage is a well-described consequence of CS exposure and growing evidence from genetic association studies and animal models of disease have suggested an important role for cellular responses to DNA damage in the pathobiology of COPD (4–8). DNA damage occurs in all cells from endogenous (e.g. metabolism) and exogenous (e.g. CS) sources. To promote genomic stability, cells maintain a network of intertwined signaling pathways collectively referred to as the DNA damage response (DDR) (9). The DDR coordinates cell-cycle checkpoints, DNA repair, and DNA tolerance pathways. In the setting of severe DNA damage, the DDR activates specific programs such as cellular senescence or apoptosis. However, the degree of DNA damage is not the only determinant of cell fate following DDR activation. Both the capacity to repair DNA and the propensity of cells to undergo apoptosis vary across cell/tissue types, between individuals, and can change with age or disease states (10). In COPD, there is increased activation of the DDR and pathologic consequences of DDR (e.g. apoptosis), even when compared to smokers without COPD (5, 11). While cellular mechanisms that regulate the DDR are likely to be important mediators of COPD susceptibility, there is little understanding of the specific DDR regulators that contribute to COPD pathogenesis.

MicroRNAs are short non-coding RNAs that function as epigenetic regulators of many cellular stress responses including the DDR (12). In this study, we sought to identify regulators of cellular stress responses that increase susceptibility to COPD and hypothesized that microRNAs regulating the DDR are critical determinant of COPD susceptibility. We analyzed microRNA expression in human lung tissue samples and found miR-24-3p inversely correlated with multiple measurements of disease severity in COPD. To determine the biological relevance of decreased miR-24-3p expression in COPD, we used cell culture and mouse models of CS exposure. We found inhibition of miR-24-3 increased susceptibility to CS-induced apoptosis and emphysema in mice and we found miR-24-3p suppresses the DDR via inhibition of the pro-apoptotic BH3-only protein BCL2L11 (commonly called BIM) and the homologous recombination protein BRCA1. Finally, we measured BIM and BRCA1 mRNA and protein in human lung tissue to determine if miR-24-3p inhibition of BIM and BRCA1 was relevant to human disease. We found BIM and BRCA1 are increased in lung tissue samples from subjects with COPD and both BIM and BRCA1 inversely correlated with miR-24-3p expression.

## RESULTS

### MiR-24-3p is decreased in COPD

We analyzed microRNA and mRNA microarray expression profiles of 172 lung tissue samples performed by the Lung Genomics Research Consortium (LGRC) (13–15), focusing on subjects with and without COPD and excluding subjects with a pathologic diagnosis of pulmonary fibrosis. Table 1 summarizes demographic and clinical characteristics of these 172 subjects. 17 microRNAs positively correlated with FEV_1_ percent predicted and 6 negatively correlated with FEV_1_ percent predicted (FDR<0.05) (Figure 1A). Of 23 correlated microRNAs, 3 microRNAs negatively correlated with percent radiographic emphysema: miR-181d-3p (ρ=-0.346), miR-551b-3p (ρ=-0.347), and miR-24-3p (ρ=-0.353). All microRNAs correlated with COPD severity measurements are shown in **Supplemental Table 1**.

**Table 1.** LGRC = Lung Genomics Research Consortium, GOLD = Global initiative for Obstructive Lung Disease, W=White, H=Hispanic, AA=African American, A=Asian, O=other, F=former smokers, N=never smokers, C=current smokers. [] represents missing data. () represents interquartile range.

|  | LGRC |  |  |
| --- | --- | --- | --- |
|  | No COPD | GOLD I, II | GOLD III, IV |
| Subjects | 78 | 50 | 44 |
| Female sex (%) | 45 (58%) | 24 (48%) | 25 (57%) |
| Age | 65 (59 - 73) | 69 (62 - 74) | 63 (54 - 69) |
| Race or ethnic group | 73 W, 1 H, 2 AA, 1 A, 1 O | 48 W, 1 AA | 42 W, 1 AA, 1 O |
| FEV<sub>1</sub> percent predicted | 94 (86 - 103) | 61 (53 - 73) [1] | 23 (19 - 33) [1] |
| % radiographic emphysema | 0.24 (0.13 - 0.52) [48] | 3.35 (0.79 - 10.88) | 25.80 (15.95 - 32.45) [2] |
| smoking status | 46 F, 22 N, 1 C [9] | 45 F, 1 N, 4 C | 41 F, 1 N, 2 C |
| Pack-Year | 10 (0 - 45.50) [13] | 50 (38 - 80) | 44 (29 - 70) [1] |

**Figure 1.**
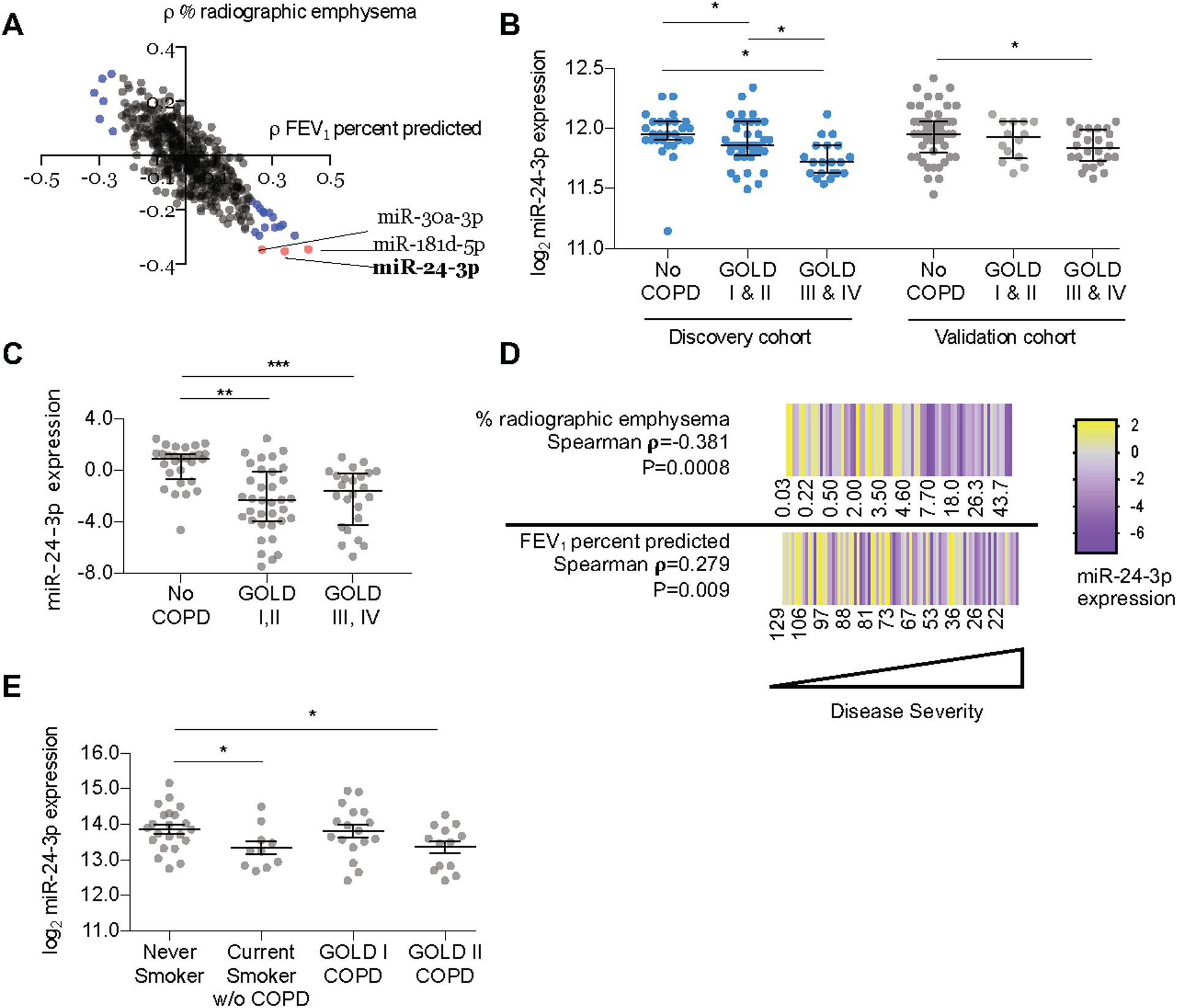
MiR-24-3p is decreased in COPD and inversely correlates with disease severity. **A)** Coefficients of Spearman correlations (ρ) between microRNAs vs. percent radiographic emphysema (y-axis) (n=121) and microRNAs vs. FEV_1_ percent predicted (*x-axis*) (n=172) in the LGRC cohort. Blue = microRNAs correlated with FEV_1_ percent predicted (FDR<0.05). Red = microRNAs correlated with FEV_1_ and percent radiographic emphysema (FDR<0.05). **B)** Log_2_-transformed microarray expression of miR-24-3p in the discovery and validation LGRC cohorts. Discovery cohort: n=28 for No COPD, n=36 for GOLD I&II, n=20 for GOLD III&IV. Validation cohort: n=50 for No COPD, n=14 for GOLD I&II, n=24 for GOLD III&IV. **C)** miR-24-3p expression (∆Ct miR-24-3p/RNU48) measured by RT-PCR in lung tissue samples from the confirmatory cohort. n=28 for No COPD, n=35 for GOLD I/II COPD, and n=24 for GOLD III,IV COPD. **D)** Heatmap of miR-24-3p expression (∆Ct miR-24-3p/RNU48) measured by RT-PCR in lung tissue samples from the confirmatory cohort vs. FEV_1_ percent predicted (n=87) and percent radiographic emphysema (n=75) (Spearman **ρ**). Yellow denotes increase above the sample median, and purple denotes decrease from the sample median. **E)** Log_2_-transformed microarray expression of miR-24-3p in airway brushings from the COSMIC cohort. n=22 for never smokers, n=10 for current smokers without COPD, n=17 for current and former smokers with COPD (GOLD I), n=13 for current and former smokers with COPD (GOLD II). Error bars represent median ± interquartile range (**B,C**) or mean ± SEM (**E**). ***P ≤ 0.0001, **P<0.001, *P<0.05, Kruskal-Wallis 1-way ANOVA (**B,C**) or ordinary one-way ANOVA (**E**), correcting for multiple comparisons using 2-stage linear step-up procedure of Benjamini, Krieger, and Yekutieli.

We focused on miR-24-3p because miR-24-3p best correlated with radiographic emphysema and miR-24-3p is highly expressed in the lung (16, 17). We then sought to validate our findings by assessing miR-24-3p expression in multiple cohorts. The LGRC cohort was previously divided into discovery and validation cohorts (**Supplemental Table 2)** and we identified decreased miR-24-3p in subjects with severe COPD in both cohorts (Figure 1B). We then measured miR-24-3p by RT-PCR in 92 lung tissue samples from subjects in an additional confirmatory cohort. Demographic and clinical characteristics of subjects in this confirmatory cohort are shown in **Supplemental Table 3**. We found decreased miR-24-3p in patients with Global Initiative for Chronic Obstructive Lung Disease (GOLD) I/II disease (0.36-fold, P<0.0001) and GOLD III/IV (0.27-fold, P<0.0001) (Figure 1C). We also found miR-24-3p expression positively correlated with FEV_1_ percent predicted (ρ=0.279, P=0.009) and negatively correlated with percent radiographic emphysema (ρ=-0.381, P=0.0008) (Figure 1D).

We then sought to validate our findings in an independent cohort with different subject characteristics and sampling techniques, so we turned to the Karolinska COSMIC study (Clinical & Systems Medicine Investigations of Smoking-related Chronic Obstructive Pulmonary Disease), a cross-sectional study in which microarrays were performed on epithelial brushings (18). Compared to subjects in the LGRC, COSMIC has a higher proportion of active smokers and subjects with less severe disease as summarized in **Supplemental Table 4**. We found miR-24-3p was decreased in epithelial brushings from active smokers without COPD (0.69-fold, P=0.031) and brushings from current and former smokers with GOLD II COPD (0.70-fold, P=0.033) (Figure 1E). Finally, we took note of a previous study by Ezzie *et al*, in which miR-24-3p was amongst the top 10 microRNAs decreased in lung tissue samples from subjects with COPD (19). While miR-24-3p was not the focus of that study, taken together with the above findings, these data from multiple cohorts provide substantial evidence that miR-24-3p is decreased in COPD and inversely correlates with COPD severity.

### Inhibition of miR-24-3p increases CS-induced apoptosis and emphysema in mice

To determine the role of miR-24-3p in human COPD, we inhibited miR-24-3p in AKR/J mice using a locked nucleic acid (LNA)-inhibitor and exposed mice to 20 weeks of CS. Study design is shown (Figure 2A). First, we demonstrated effective inhibition of miR-24-3p 8 weeks (0.061-fold) and 10 weeks (0.62-fold) after intranasal delivery of the miR-24-3p LNA inhibitor (Figure 2B). Mice were then treated with miR-24-3p LNA-inhibitor or LNA-control and exposed to CS for 20 weeks, which is the approximate length of time required for the development of emphysema in AKR/J mice (20). As expected, mice exposed to CS demonstrated increased lung compliance, airspace enlargement, and apoptosis. Impressively, miR-24-3p inhibition significantly enhanced CS-mediated increases in lung compliance (20.3% vs. 14.8%) (Figures 2C,D), air space enlargement as measured by mean linear intercept (24.8% vs. 15.2%) (Figures 2E,F) and apoptosis (1.84-fold) (Figure 2G, **Supplemental Figure 1**). These results demonstrate miR-24-3p inhibition enhances susceptibility to apoptosis and emphysema in a CS-exposure murine model.

**Figure 2.**
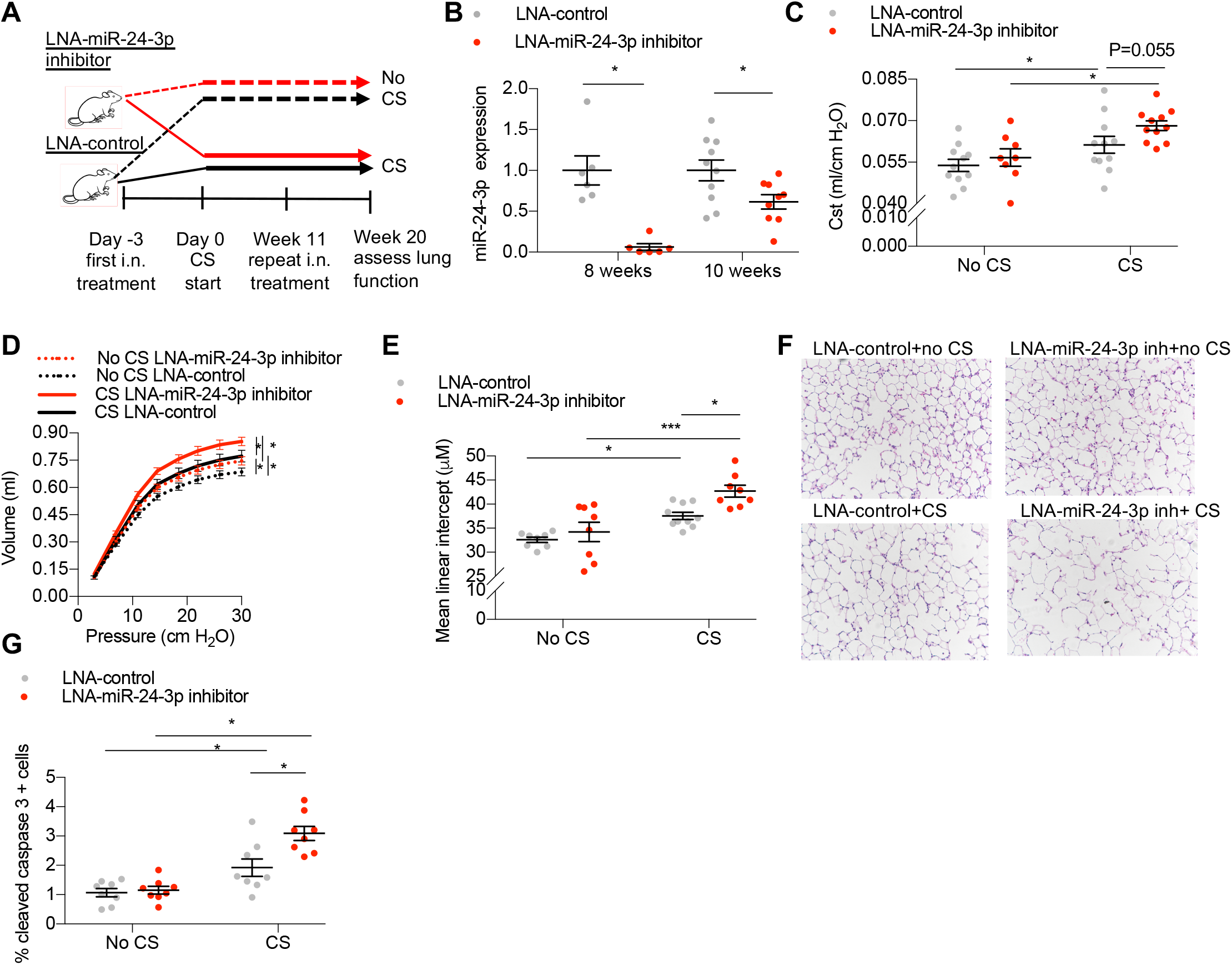
Inhibition of miR-24-3p increases susceptibility to CS-induced apoptosis and emphysema. **A)** Study design for mice treated with intranasal (i.n.) LNA-miR-24-3p inhibitor or LNA-control ± cigarette smoke (CS). **B)** Relative expression of miR-24-3p (∆∆Ct miR-24-3p/snoRNA202) measured by RT-PCR in mouse lungs at 8 or 10 weeks following intranasal administration (i.n.) with locked nucleic acid (LNA) miR-24-3p inhibitor or LNA-control. n=6-10/group. **(C-F)** Lung function and histologic assessments of emphysema in mice exposed to CS (n=11/LNA-control and n=11/LNA-miR-24-3p inhibitor) and mice exposed to no CS (n=13/LNA-control and n=8/LNA-miR-24-3p inhibitor). **C)** Flexivent measurements of static lung compliance. **D)** Lung compliance assessed by the slope of the pressure-volume deflation limb. **E-F**) Mean linear intercept (***µ***M) in mice treated with LNA-miR-24-3p inhibitor vs. LNA control ± CS with representative hematoxylin and eosin lung histology. Images acquired with 20x objective. **G)** Apoptosis detected by quantification of cleaved caspase-3 in lungs from mice treated with LNA-miR-24-3p inhibitor vs. LNA-control ± CS exposure. n=8/group. Error bars mean ± SEM. ***P ≤ 0.0001, **P<0.001, *P<0.05, ordinary one-way ANOVA correcting for multiple comparisons using 2-stage linear step-up procedure of Benjamini, Krieger, and Yekutieli.

### MiR-24-3p inhibits apoptosis by targeting BIM

We then sought to determine the role of miR-24-3p in epithelial cell responses to CS. To modulate miR-24-3p expression, we used miR-24-3p mimics and miR-24-3p inhibitors to overexpress (6.4-fold) and inhibit (0.039-fold) miR-24-3p respectively (**Supplemental Figure 2**). Cigarette smoke extract (CSE) was used to model epithelial response to CS. Transfection of human airway epithelial cells (HAECs) with miR-24-3p inhibitor increased susceptibility to CSE-mediated apoptosis (1.27-fold), while transfection with miR-24-3p mimic decreased susceptibility to CSE-mediated apoptosis (0.19-fold) (Figure 3A). MiR-24-3p also inhibited CSE-mediated apoptosis in BEAS-2B cells as assessed by flow cytometric measurements of Annexin/PI (0.32-fold) and caspase-3/7 (0.62-fold) (**Supplemental Figure 3**).

**Figure 3.**
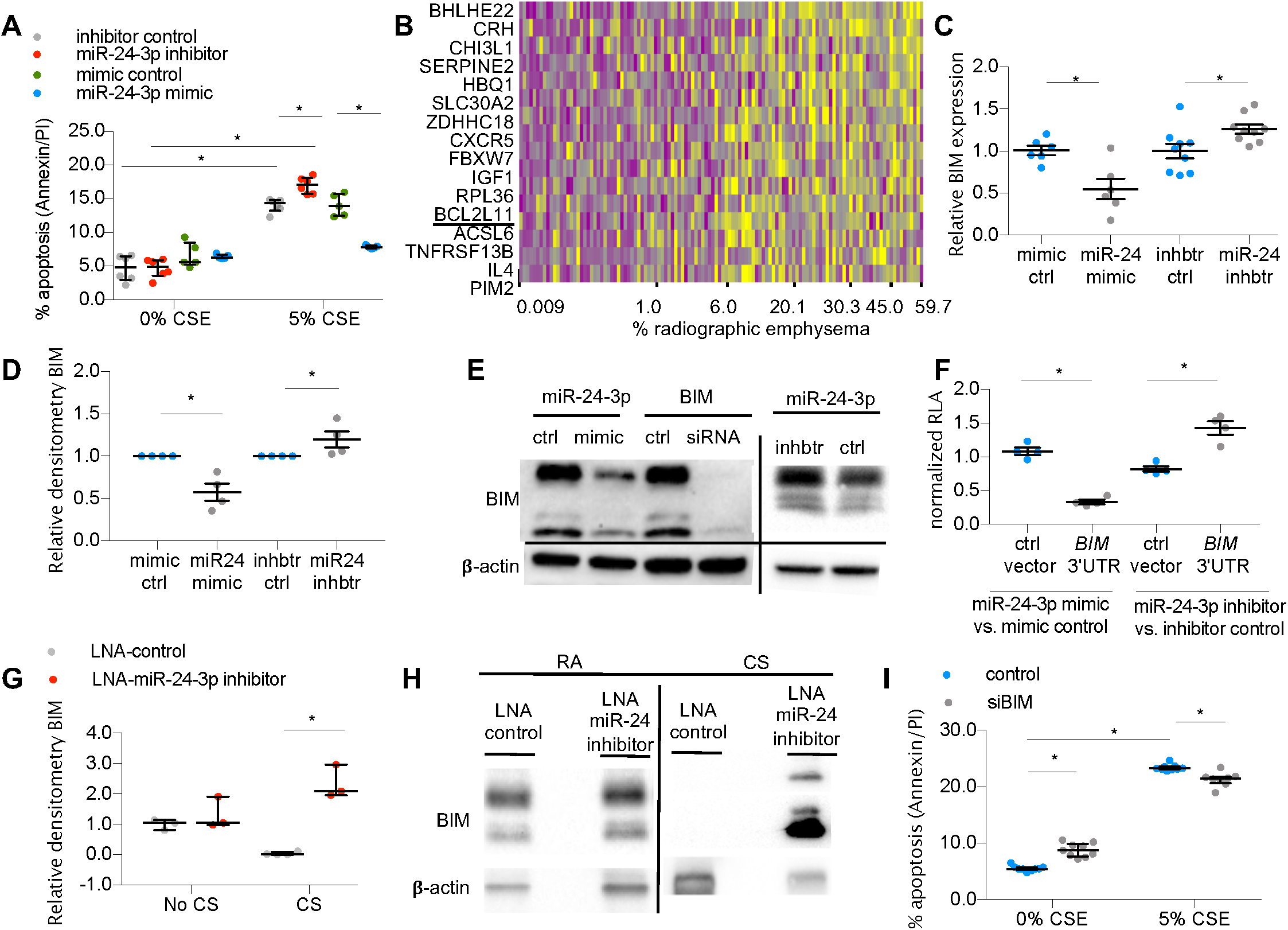
MiR-24-3p inhibits apoptosis through BIM. **A**) Percent apoptotic cells determined by flow cytometry for Annexin V/propidium iodide (PI) in miR-24-3p mimic vs mimic control (n=5/group), and miR-24-3p inhibitor vs. inhibitor control (n=6/group) treated human airway epithelial cells exposed to 0% or 5% cigarette smoke extract (CSE). **B)** Heatmap of z-scores of miR-24-3p target genes, as measured by microarray expression in the LGRC cohort correlated with percent radiographic emphysema (Spearman **ρ**, FDR<0.05). n=121 subjects. Yellow denotes increase above sample median, and purple denotes decrease from sample median. **C)** BIM expression (∆∆Ct BIM/18s) measured by RT-PCR in BEAS-2B cells treated with miR-24-3p mimic vs mimic control (n=6/group), and miR-24-3p inhibitor vs. inhibitor control (n=9/group). **D-E**) Relative densitometry of BIM/β;-actin in BEAS-2B cells treated with miR-24-3p mimic vs. mimic control or miR-24-3p inhibitor and miR-24-3p inhibitor vs. inhibitor control. n=4/group. Sample immunoblotting includes siRNA against BIM and siRNA control. **F)** Relative luciferase activity (firefly luciferase/Renilla luciferase) normalized as a ratio of miR-24-3p mimic vs. mimic control in BEAS-2B or miR-24-3p inhibitor vs. inhibitor control cells co-transfected with BIM 3’UTR luciferase reporter plasmid or control plasmid (n=4/group). **G-H**) Relative densitometry of BIM/β-actin from immunoblots performed on lung tissue lysates from mice treated with LNA-control or LNA-miR-24-3p inhibitor exposed to 20 weeks of cigarette smoke (CS) (n=4/group) or no CS (n=3/group) with sample immunoblotting shown. **I)** Percent apoptotic cells determined by flow cytometry for Annexin V/PI in BEAS-2B cells treated with siBIM or siRNA control and exposed to 0% or 5% CSE (n=9/group). Error bars represent median ± interquartile range. *P<0.05, Kruskal-Wallis correcting for multiple comparisons using 2-stage linear step-up procedure of Benjamini, Krieger, and Yekutieli.

To determine the mechanism through which miR-24-3p inhibited apoptosis, we started by immunoblotting for p53 and phosphorylated p53 (Ser15) and found miR-24-3p inhibition of CSE-mediated apoptosis was p53-independent (**Supplemental Figure 4**). We then sought to determine the p53-independent mechanism via which miR-24-3p inhibits apoptosis. Therefore, we reanalyzed the 172 patient LGRC dataset to identify miR-24-3p target genes correlated with disease severity. We identified 1417 genes inversely correlated with miR-24-3p expression, of which 57 were putative miR-24-3p targets as predicted by TargetScan, miRDB, or mirTarbase. Of these 57 genes, 16 genes positively correlated with percent radiographic emphysema (FDR<0.05) (Figure 3B) including BCL2L11 (BIM), a pro-apoptotic BH3-only protein (21). In order to determine if miR-24-3p regulates BIM in airway epithelial cells, we transfected BEAS2B cells with miR-24-3p mimic, miR-24-3p inhibitor, and respective controls. MiR-24-3p mimic decreased BIM mRNA (0.548-fold) and protein (0.575-fold), while miR-24-3p inhibitor increased BIM mRNA (1.26-fold) and protein (1.20-fold) (Figures 3C-E). Notably, miR-24-3p decreased all three isoforms of BIM (BIM EL, BIM L, and BIM S) (Figure 3E). We transfected BEAS2B cells with an expression vector containing the 3’untranslated region (3’UTR) of *BIM* upstream of the firefly luciferase gene and demonstrated that miR-24-3p targets the 3’UTR of *BIM* for degradation (Figure 3F). We confirmed miR-24-3p modulates BIM *in vivo* by immunoblotting for BIM in lung tissue lysates from mouse studies and finding increased BIM in mice treated with LNA-miR-24-3p inhibitor and exposed to CS (63.6-fold) (Figures 3G,H). Finally, we sought to determine the role of BIM in CSE-mediated apoptosis in epithelial cells. After demonstrating effective inhibition of BIM with siRNA (siBIM) (**Supplemental Figure 5**), we found siBIM decreased CSE-mediated apoptosis in BEAS-2B cells as assessed by flow cytometric measurements of Annexin/PI (0.91-fold) and caspase 3/7 (0.86-fold) (Figure 3I, **Supplemental Figure 6**), albeit to a lesser extent than transfection with miR-24-3p mimic. These data suggest miR-24-3p inhibits apoptosis, in part, through BIM.

### MiR-24-3p impairs double stranded DNA break repair by homologous recombination through BRCA1

To understand miR-24-3p inhibition of apoptosis within the larger context of the DDR, we sought to determine the effects of miR-24-3p on DNA repair. Previous gene ontology enrichment analyses suggested genes involved in the repair of double stranded DNA breaks (DSBs) are overrepresented amongst miR-24-3p target genes (22). DSBs are deleterious types of DNA damage caused by CS (**Supplemental Figure 7**). Cells respond to DSBs by phosphorylating histone H2AX (γH2AX) and recruiting DNA repair proteins such as p53-binding protein 1 (53BP1) to form γH2AX foci and colocalized γH2AX/53BP1 foci (9). To model the role of miR-24-3p in DSB repair, we used ionizing radiation, a well-known inducer of DSBs, and imaging-flow cytometry for high-throughput and automated detection of DNA repair foci (Figure 4A)(23). In HAECs, transfection with miR-24-3p mimic inhibited resolution of γH2AX foci and colocalized γH2AX/53BP1 foci following IR, suggesting miR-24-3p inhibits DSB repair (Figures 4B,C). We measured DAPI intensity to estimate cell cycle phase (G_1_ and S/G_2_) and observed no differences in cell cycle phase between HAECs treated with mimic control and miR-24-3p mimic (**Supplemental Figure 8**). We also observed diminished DSB repair in miR-24-3p treated cells even after dichotomizing samples by cell cycle phase (**Supplemental Figures 9 A,B**). We performed the same experiments in BEAS2B cells and similarly observed decreased DSB repair following transfection with miR-24-3p (**Supplemental Figures 9 C,D**). Gating strategy is shown (**Supplemental Figure 10**).

**Figure 4.**
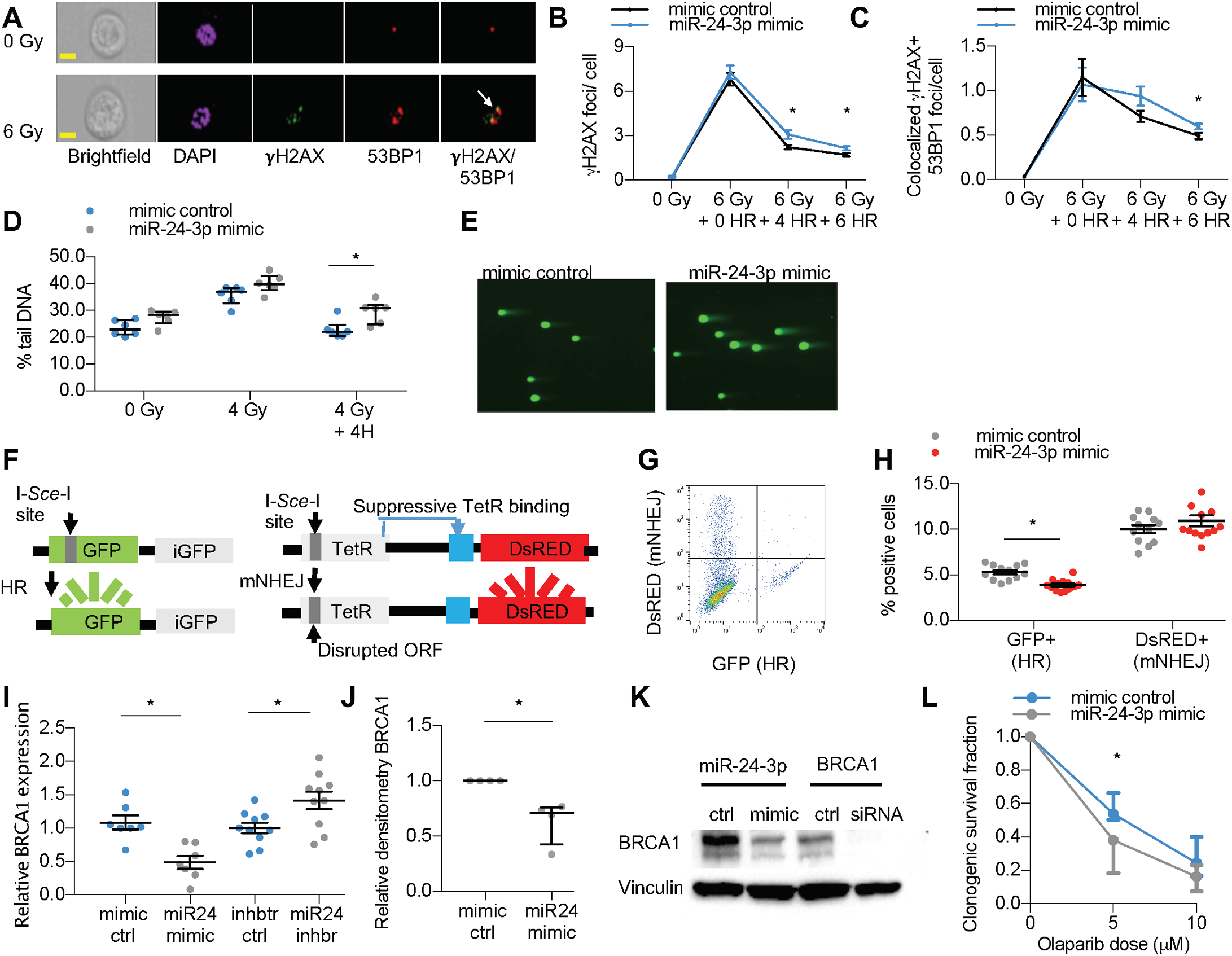
MiR-24-3p inhibits homologous recombination (HR) and BRCA1. Imaging flow cytometry of cells exposed to 0 or 6 Grey (Gy) of ionizing radiation. DAPI (blue), FITC-conjugated anti-***γ***H2AX (green), and APC-conjugated anti-53BP1 (red). Colocalized ***γ***H2AX/53BP1 foci (yellow) are shown (arrow). Yellow bar = 10µM. **B-C)** Human airway epithelial cells (HAECs) exposed to 0 Gy or 6 Gy with 0-6 hours (H) recovery. n=7/group. **D-E)** Comet assay in BEAS-2B cells exposed to 0Gy, 4Gy, or 4Gy with 4 H recovery. Percent tail DNA reflects DNA damage. n=6/group, with sample images (20x) following 4Gy with 4H of recovery. **F)** Schema of DNA reporter cell assay with 2 integrated loci for measuring homologous recombination (HR) and mutagenic non-homologous end joining (mNHEJ). **G)** Representative flow cytometry demonstrating DsRED+ (mNHEJ) and GFP+ (HR) expression. **H**) DNA reporter cells transfected with miR-24-3p mimic (n=13/group) vs. mimic control (n=12/group). **I)** BRCA1 expression (∆∆Ct of BRCA1/18s) measured by RT-PCR in BEAS-2B cells treated with miR-24-3p mimic vs. mimic control (n=7/group) and miR-24-3p inhibitor vs. inhibitor control (n=10/group). **J-K)** Relative densitometry of BRCA1/Vinculin in BEAS-2B cells treated with miR-24-3p mimic vs. mimic control. n=4/group. Sample immunoblotting includes siRNA against BRCA1 and siRNA control. **L)** BEAS-2B cells transfected with miR-24-3p mimic vs. mimic control and treated with Olaparib at indicated dosages. n=5/group. Error bars represent mean ± SEM (**B,C,H,I)** or median ± interquartile range (**D,J,L**). *P<0.05 ordinary one-way ANOVA (**B, C,H,I)**, Mann-Whitney **(J,L**) or Kruskal-Wallis **(D**) correcting for multiple comparisons using 2-stage linear step-up procedure of Benjamini, Krieger, and Yekutieli.

To confirm miR-24-3p inhibits DSB repair, we used the neutral comet assay and again found miR-24-3p impaired resolution of DSBs in BEAS2B cells (0.83-fold) (Figure 4D,E). DSB repair occurs through two canonical pathways, homologous recombination (HR) and the more error prone non-homologous end joining (NHEJ). We utilized a novel reporter cell line that distinguishes between mutagenic non-homologous end joining and homologous recombination (Figures 4F,G) and found miR-24-3p inhibited HR (0.48-fold), but not mutagenic NHEJ (Figure 4H) (24). We did not find any well-characterized HR genes that correlated with miR-24-3p expression in human lung tissue samples. However, a previous study suggested that miR-24-3p inhibits the canonical HR protein BRCA1 (22). Transfection of BEAS2B cells with miR-24-3p mimic inhibited BRCA1 expression (0.45-fold) and transfection with miR-24-3p inhibitor increased BRCA1 expression (1.42-fold) (Figure 4I). Transfection with miR-24-3p mimic also decreased BRCA1 protein (0.63-fold) (Figures 4J,K). Finally, we wanted to determine if transfection with mir-24-3p mimic induced functional BRCA1 deficiency. BRCA1 deficient cells are sensitive to poly (ADP-ribose) polymerase (PARP) inhibitors, such as Olaparib, via synthetic lethal interactions (25). As expected, transfection with miR-24-3p increased susceptibility of BEAS-2B cells to Olaparib as measured by clonogenic survival assay (Figure 4L). These data suggest miR-24-3p impairs HR by inhibiting BRCA1.

### BIM is elevated in human COPD and inversely correlates with miR-24-3p expression

To determine the relevance of our findings to human disease, we measured BIM in human lung tissue samples. RT-PCR measurement of BIM expression in the confirmatory cohort demonstrated BIM mRNA inversely correlates with miR-24-3p expression (ρ=-0.238, P=0.03) (Figure 5A). BIM mRNA is increased in patients with GOLD III,IV COPD (1.74-fold, P=0.009) (Figure 5B). BIM mRNA also inversely correlates with FEV_1_ percent predicted (ρ =-0.265, P= 0.02) and BIM mRNA correlates with percent radiographic emphysema (ρ=0.364, P=0.0001) (Figure 5C). We performed immunoblotting for BIM on lysates of frozen lung tissue samples obtained from the Lung Tissue Research Consortium (LTRC). Demographic and clinical characteristics of subjects are shown (**Supplemental Table 5**). We found increased BIM in subjects with GOLD IV COPD compared to GOLD I COPD (1.77-fold) (Figures 5D,E). Together, these data suggest BIM is increased in the lungs of patients with severe COPD and inversely correlate with miR-24-3p expression.

**Figure 5.**
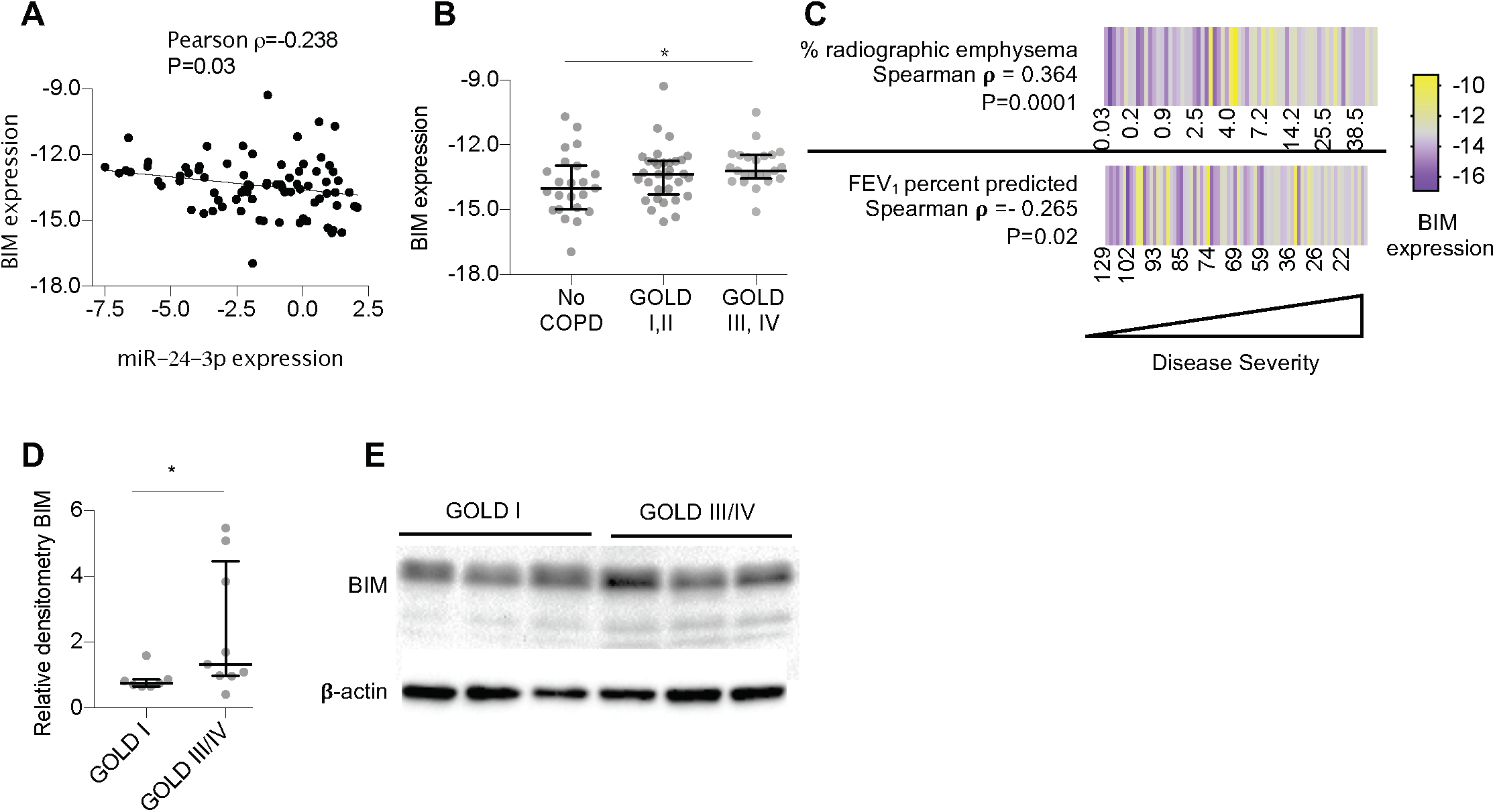
BIM inversely correlates with miR-24-3p expression and is increased in COPD. **A)** Pearson correlation of BIM expression (∆Ct BIM/18s) and miR-24-3p expression (∆Ct miR-24-3p/RNU48) measured by RT-PCR in lung tissue samples from the confirmatory cohort (n=78). **B**) BIM expression (∆Ct BIM/18s) measured by RT-PCR in lung tissue samples from the confirmatory cohort; n=23 for No COPD, n=32 for GOLD I/II COPD, and n=23 for GOLD III,IV COPD. **C)** Heatmap of FEV_1_ percent predicted (n=78) and percent radiographic emphysema (n=68) correlated with BIM expression (∆Ct BIM/18s) measured by RT-PCR in lung tissue samples from the confirmatory cohort (Spearman **ρ**). Yellow denotes increase above the sample median and purple denotes decrease from the sample median. **D-E)** Relative densitometry of BIM/β-actin from immunoblots performed on lung tissue samples from individuals with GOLD I (n=7/group) or GOLD III/IV COPD (n=9/group). Sample immunoblot is shown. Error bars represent median ± interquartile range *P<0.05 Kruskal-Wallis (**B**) or Mann-Whitney (**D**) correcting for multiple comparisons using 2-stage linear step-up procedure of Benjamini, Krieger, and Yekutieli.

### BRCA1 is elevated in human COPD and inversely correlates with miR-24-3p expression

We also measured BRCA1 in human lung tissue samples. RT-PCR measurements of BRCA1 expression in the confirmatory cohort demonstrated BRCA1 mRNA inversely correlates with miR-24-3p expression (ρ=-0.372, P=0.0008) (Figure 6A). We also found increased expression in BRCA1 with GOLD I/II COPD (3.58-fold, P=0.0001) and GOLD III/IV COPD (1.79-fold, P=0.04) (Figure 6B). Notably, we also identified a significant decrease in BRCA1 mRNA expression between GOLD I/II and GOLD III/IV disease (0.60-fold, P=0.01). We then used Automated Quantitative Analysis (AQUA) to quantify BRCA1 protein in human lung tissue. AQUA is a validated method for objectively quantifying protein within defined subcellular compartments (26). Images from lung tissue samples were obtained for three different channels: DAPI to visualize the nucleus, cytokeratin to identify epithelial cells, and BRCA1 (Figure 6C). Binarization of the DAPI and cytokeratin signal created an image mask of epithelial nuclei, within which we quantified BRCA1 signal intensity. We analyzed BRCA1 in a subset of available paraffin-embedded tissue samples available through the LTRC. Demographic and clinical characteristics of subjects are shown (**Supplemental Table 6**). First, we confirmed BRCA1 measurements by AQUA in the lung were operator independent (**Supplemental Figure 11**). We found BRCA1 inversely correlated with miR-24-3p expression (ρ=-0.813, P<0.0001) (Figure 6D) and we found increased BRCA1 in cytokeratin positive nuclei from COPD lung tissue samples compared to non-COPD samples (1.76-fold, P<0.0001) (Figure 6E). BRCA1 inversely correlated with FEV_1_ percent predicted (ρ=-0.730, P<0.0001) (Figure 6F) and BRCA1 correlated with percent radiographic emphysema (ρ=0.471, P=0.03) (Figure 6G). Together, these data suggest BRCA1 is increased in the lungs of patients with COPD and inversely correlate with miR-24-3p expression.

**Figure 6.**
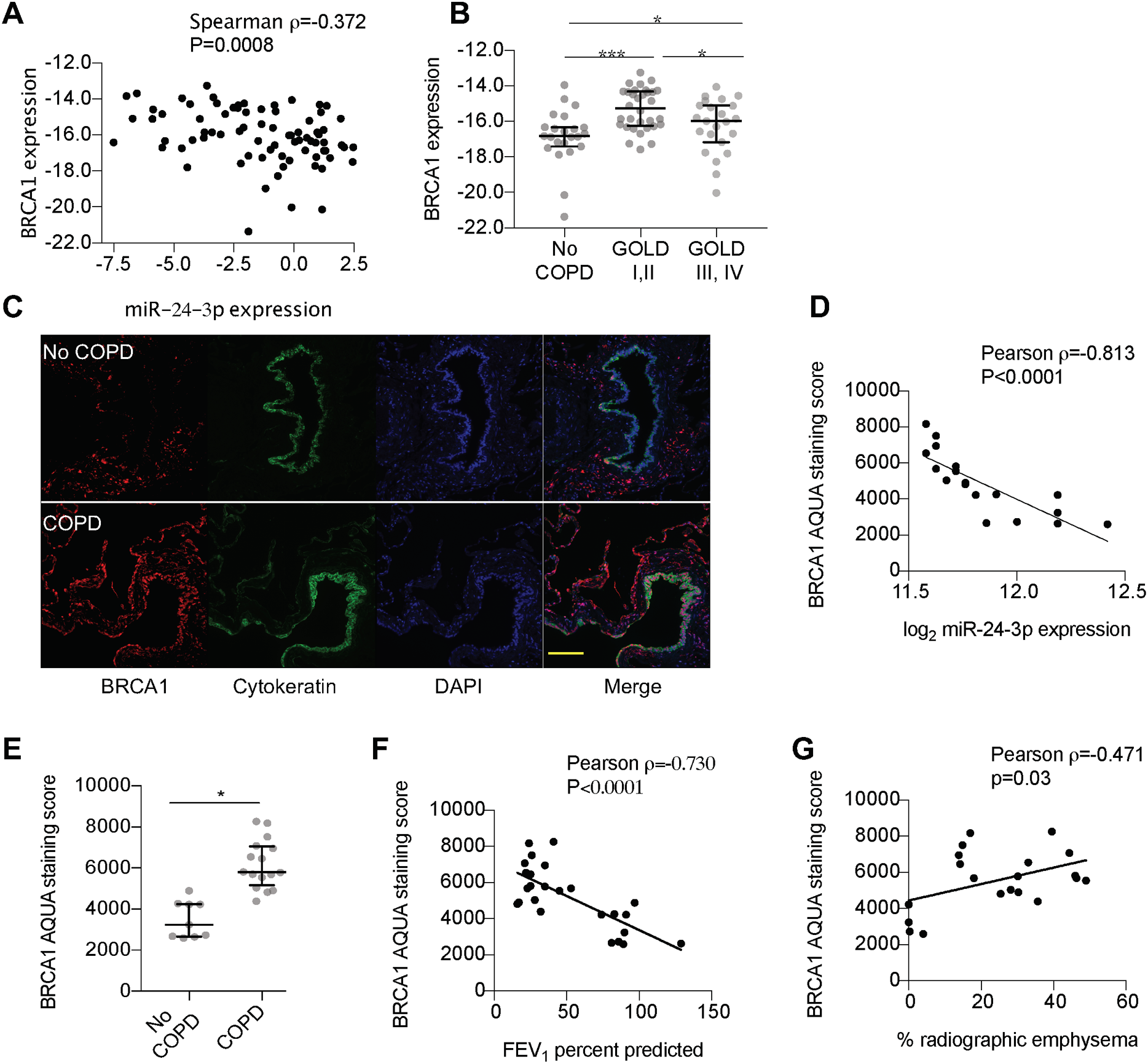
BRCA1 inversely correlates with miR-24-3p expression and is increased in COPD. Pearson correlation of BRCA1 expression (∆Ct BRCA1/18s) and miR-24-3p expression (∆Ct miR-24-3p/RNU48) measured by RT-PCR in lung tissue samples from the confirmatory cohort (n=78). **B**) BRCA1 expression (∆Ct BRCA1/18s) measured by RT-PCR in lung tissue samples from the confirmatory cohort; n=23 for No COPD, n=32 for GOLD I/II COPD, and n=23 for GOLD III,IV COPD. **C)** Representative fluorescence microphotographs showing in situ detection of BRCA1, cytokeratin, and DAPI nuclear stain. BRCA1 staining intensity within the image mask generated from the cytokeratin and DAPI positive staining was used to generate a quantitative score of BRCA1 staining using Automated Quantitative Analysis (AQUA). Yellow bar=50 µM **D)** Pearson correlation between miR-24-3p expression and BRCA1 AQUA staining scores (n=19). **E)** BRCA1 AQUA staining scores from control (n=9/group) and COPD subjects (n=16/group). **F)** Pearson correlation between FEV_1_ percent predicted and BRCA1 AQUA staining scores (n=25). **G)** Pearson correlation between percent radiographic emphysema and BRCA1 AQUA staining scores (n=21). Error bars represent median ± interquartile range (**B**) or mean ± SEM (**E**). *P<0.05 Kruskal-Wallis (**B**) correcting for multiple comparisons using 2-stage linear step-up procedure of Benjamini, Krieger, and Yekutieli or student t-test (**E**).

## DISCUSSION

In this study, we analyzed microRNA profiles of lung tissue samples from subjects with and without COPD and identified that miR-24-3p was decreased in COPD and inversely correlated with disease severity (FEV_1_ percent predicted and radiographic emphysema). Decreased miR-24-3p increases susceptibility to CS-mediated emphysema as demonstrated in a CS exposure-murine model. We also demonstrated that miR-24-3p inhibits apoptosis and homologous recombination, and that these effects were partially mediated through inhibition of BIM and BRCA1 respectively. Finally, we found that BIM and BRCA1 are increased in patients with COPD and inversely correlate with miR-24-3p expression. Together, these findings suggest that decreased miR-24-3p primes the DDR and contributes to disease progression in COPD.

We identified multiple microRNAs that correlated with continuous measurements of airflow obstruction and emphysema (nominal P<0.05) including COPD-associated microRNAs such as: miR-218-5p(27), miR-126-3p(8), miR-223-3p(19), miR-199a-5p(28) and multiple members of the miR-181, miR-34, and miR-30 microRNA families(29–31) (**Supplemental Table 1**). However, the microRNA that best correlated with radiographic emphysema was miR-24-3p. MiR-24-3p is a member of the miR-23-27-24 microRNA family which is found in two genomic loci in humans(22). While the expression of miR-27a and miR-23a correlated with FEV_1_ percent predicted and percent radiographic emphysema, the expression of miR-27b and miR-23b did not. Therefore, decreased miR-24-3p expression may be related to the function of the miR-23-27-24 cluster on chromosome 19p13. Notably, miR-181d is also located within 19p13 and miR-181d was one of the top 3 microRNAs that correlated with radiographic emphysema in our study (Figure 1A). Therefore, this chromosomal region may be important in mediating disease pathogenesis and future studies are warranted.

We found that miR-24-3p inhibits apoptosis in part through the pro-apoptotic BH3-only protein BIM and BIM is increased in the lungs of patients with COPD. Apoptosis is a key mechanism of COPD pathobiology and required for the development of emphysema in mice (11). The propensity of cells to undergo apoptosis in response to injury (i.e. “apoptosis priming”) varies across tissue and cell types (32). BIM has been demonstrated to promote apoptosis under physiologic and pathophysiologic conditions (21). While BIM is regulated by diverse signaling mechanisms, post-transcriptional regulation of BIM by miR-24-3p contributes to disease pathology in other organ systems, such as ischemic heart disease (33). Very little is known about the role of BIM in COPD, although increased BIM was identified in a murine model of copper deficiency-induced emphysema (34). Our data suggest that decreased miR-24-3p inhibits BIM and increases cellular susceptibility to apoptosis and disease severity in COPD.

Our data also shows that miR-24-3p suppresses HR repair of DSBs. Previous studies have also shown that mIR-24-3p inhibits DNA repair, but these studies have primarily focused on miR-24-3p suppression of H2AX (35). While H2AX may indeed be a target of miR-24-3p, we did not observe reduced foci of H2AX phosphorylation (i.e. γH2AX foci) or diminished mutagenic NHEJ following treatment of cells with miR-24-3p mimics. However, we did identify a role for miR-24-3p inhibition of HR and the canonical HR protein BRCA1. We also found that BRCA1 mRNA and protein were increased in COPD lung tissue and inversely correlated with miR-24-3p. This up-regulation of BRCA1 may seem paradoxical at first. BRCA1 is involved in DNA repair and has anti-oxidant effects (36). However, the overexpression of single DNA repair proteins can dysregulate complex DNA repair steps (37). BRCA1 can also facilitate stress-induced apoptosis and activate inflammatory GADD45a signaling which may contribute to COPD pathogenesis (38, 39). An increase in epithelial expression of BRCA1 is a novel finding in COPD and suggests that decreased miR-24-3p does not simply increase apoptosis priming but priming of the whole DDR. This notion that regulatory elements prime the DDR in COPD is supported by a recent letter by Paschalaki *et al* who identified that miR-126-3p is decreased in COPD and miR-126-3p suppresses the DDR protein ataxia telangiectasia mutated (ATM)(8). Future studies will be necessary to determine the role of specific DDR elements, such as BRCA1, in COPD pathogenesis.

A key strength of this study is the large number of subjects in the LGRC cohort which allowed us to identify microRNAs that *correlated* with COPD relevant features. Categorical definitions of disease severity (e.g. GOLD stage) do not fully address interpatient variability and can both over- and underestimate disease severity (40). Many have suggested that a better approach to understanding COPD pathogenesis is through the evaluation of multiple continuous disease traits which our large sample size allowed us to do (2, 41). However, there are several limitations to this work that deserve mention. First, this is a cross-sectional study and it is difficult to confirm that decreased miR-24-3p contributed to COPD pathogenesis in humans. To address this concern, we used an animal model of COPD to demonstrate proof-of-principle that decreased miR-24-3p increases susceptibility to CS-mediated emphysema. A second limitation is that we profiled whole lung tissue samples and therefore differential gene expression may be due to varying proportions of lung cell types. However, miR-24-3p is not specifically expressed in a single cell type (17). Additionally, we found decreased miR-24-3p specifically in airway epithelial cell brushings from patients with COPD in the COSMIC cohort and we identified up-regulation of BRCA1 specifically in cytokeratin positive epithelial cells. A third limitation is that microRNAs can target many different proteins and pathways, therefore, the DDR may not be the only miR-24-3p target in COPD pathogenesis. For example, miR-24-3p was shown to inhibit T-cell production of IL-4 production and therefore miR-24-3p expression may also modulate airway inflammation to increase susceptibility to COPD (42). Further studies will be necessary to confirm the role of downstream miR-24-3p targets in mediating COPD pathogenesis. Finally, there is inherent selection bias when obtaining tissue samples from COPD patients. Subjects with early stage or no COPD are likely being evaluated for a radiographic finding concerning for malignancy while samples from subjects with severe disease are commonly taken from explanted lungs. Even though lung tissue samples are obtained at a distance from suspicious lesions, there is always a concern for a “field of cancerization” effect and the potential role for malignancy as a confounding variable (43). However, a previous array-based study of lung tissue from subjects without cancer also demonstrated decreased miR-24-3p expression in COPD (19).

A key implication of our finding is that disease progression in COPD may involve exaggerated cellular stress responses to DNA damage as a consequence of microRNA expression changes. This exaggerated DDR may drive pathologic processes like apoptosis regardless of *functional* DNA repair capacity. Developing a more robust understanding of the various arms of the DDR and the biologic mechanisms through which a dysregulated DDR contributes to the pathogenesis of COPD may lead to the development of therapies that minimize pathogenic DDR responses while preserving DNA repair capacity and ultimately provide relief for those with individuals suffering from this disease.

## METHODS

### Lung Genomics Research Consortium (LGRC) cohort

We analyzed microRNA and mRNA microarray profiles of flash frozen lung tissues samples obtained from the National Heart, Lung, and Blood Institute (NHLBI)– sponsored Lung Tissue Research Consortium (LTRC). Lung tissue samples were obtained from explanted or resected, or biopsied lung tissue. Methods of tissue procurement, cohort characteristics, and gene expression profiling have been previously described and further details are provided in Supplementary Methods (13–15). We identified 172 paired mRNA and microRNA expression profiles from patients in the LGRC cohort that did not have a pathologic diagnosis of interstitial lung disease and were designated as “control” or “COPD”. As part of the original study design, samples were previously stratified into discovery and validation cohorts.

### Confirmatory cohort for RT-PCR

We measured miR-24-3p by RT-PCR in 92 available human lung tissue samples obtained from the LTRC that that did not have a pathologic diagnosis of interstitial lung disease and were designated as “control” or “COPD.” Samples were made available through the LGRC. The TaqMan MicroRNA Reverse Transcription Kit (Thermo Scientific) was used to measure miR-24-3p and RNU48 (internal control) per the manufacturer’s protocol. All samples were run in duplicate. 5 miR-24-3p measurements were excluded due to inadequate RNA concentrations or a coefficient of variation > 7.5%. Of the remaining 87 samples, 36 subjects overlapped with the above microarray analysis while 51 subjects were unique. Subsequently, we measured BIM and BRCA1 by RT-PCR in the same samples. 18s was used as an internal control gene. Applying the same exclusion criteria, 10 additional BIM and BRCA1 samples were excluded.

### Clinical & Systems Medicine Investigations of Smoking-related Chronic Obstructive Pulmonary Disease (COSMIC) cohort

The Karolinska COSMIC cohort (Clinical Trials ID: NCT02627872) is a cross-sectional study designed for investigating the molecular pathogenesis of COPD. Methods of tissue procurement, cohort characteristics, and gene expression profiling have been previously described and further details are provided in Supplementary Methods (18, 44). MicroRNA profiling was performed on airway epithelial brushings collected by fiberoptic bronchoscopy in a subset of the Karolinska COSMIC cohort. Subjects included never-smokers, “healthy” smokers with normal lung function, and individuals with COPD (GOLD stage I–II). Smokers were matched by smoking history (>10 pack years) and current smoking habits (10 cigarettes/day the past 6 months).

### Human lung tissue samples for measuring BIM and BRCA1

Lung tissue samples used for western blot to assess BIM and paraffinized lung tissue samples for assessing BRCA1 AQUA score were obtained through the LTRC. Immunoblotting methods for measuring BIM are described in Supplemental Methods. AQUA is described below.

### Cell culture and transfection

BEAS-2B cells (American Type Culture Collection) were cultured in RPMI 1640 supplemented with 10% FBS and passaged less than 30 times. Primary human airway epithelial cells (HAECs) (Lonza) were cultured in PneumaCult-expansion medium (Stem Cell) and used through passage 6. MiR-24-3p mimic, mimic control, miR-24-3p inhibitor, inhibitor control, siRNA duplex for silencing BRCA1 (siBRCA1), and silencing RNA non-targeting control were obtained from Thermo Scientific. RNA duplexes for silencing BIM (siBIM) and non-targeting control were obtained from Dharmacon. Cells were transfected in 6-well plates using RNAiMAX transfection reagent (Life Technologies) and OptiMEM media according to manufacturer’s protocols. Cells were harvested or used for experimentation 24-48 hours after transfection. RT-PCR and western blot techniques are described in Supplemental Methods.

### Cigarette Smoke Extract (CSE)

Mainstream smoke from one 3RF4 research cigarettes (University of Kentucky) was bubbled via negative pressure through 10 mL of cell culture media in a fume hood for about 5 minutes. CSE was filtered via 0.22 µM filter (MilliporeSigma). The obtained filtrate was considered as 100% CSE. CSE was sterile filtrated, aliquoted, and stored at −80°C. CSE was quickly thawed for usage and diluted in cell-type specific media at indicated concentrations.

### Apoptosis assays

Fluorescence-activated cell sorting (FACS) analysis of transfected BEAS-2B cells for Annexin V and propidium iodide (PI) was performed per the manufacturer’s protocol (BD Biosciences). FACS analysis of transfected BEAS-2B cells for caspase 3 & caspase 7 along with SYTOX AADvanced^TM^ Dead Cell Stain was performed per manufacturer’s protocol (Thermo Scientific).

### 3’untranslated region (UTR) luciferase assay

Cells were transfected with the firefly luciferase-expressing (pEZX-MT05) plasmids containing the 3′UTR of *BIM* or an empty vector (Genecopoeia) and were co-transfected with miR-24-3p mimic, mimic control, miR-24-3p inhibitor, or inhibitor control using Lipofectamine 2000 (Invitrogen). After 24-48 hours, the cells were harvested, lysed and analyzed for firefly and Renilla luciferase using Luc-Pair Duo-Luciferase HS Assay Kit expression as described in the manufacturer’s protocol (Genecopoeia).

### AMNIS flow cytometry

Full details and gating strategy are described in Supplemental Methods. Briefly, cells were washed in permeabilization buffer (eBioscience) and stained overnight at 4ºC with FITC conjugated anti-phospho-histone H2AX (serine139) (1:2,000, Millipore) and AlexaFluoro-647 conjugated 53BP1 (1:400, Novusbio). This was followed by staining with DAPI (1:5,000) for 5 minutes. Cells were then analyzed with AMNIS imaging flow-cytometer (MilliporeSigma). At least 1000 cells were captured per condition at 40x magnification with extended depth of field. Data was analyzed using IDEAS 6.2 imaging flow cytometry software (MilliporeSigma) (23).

### Neutral Comet Assays

Neutral comet assays were performed per manufacturer’s protocol (Trevigen). Briefly, cells were trypsinized and washed with PBS. Samples were suspended in LM Agarose (Trevigen). Neutral electrophoresis was conducted at 21 V for 1 hour. Cells were stained with SYTOX Green nucleic acid stain (Thermo Scientific) and images of at least 75 cells were captured with a with a Nikon Eclipse Ti microscope.

### Double stranded break (DSB) reporter assay

The construction and details of the U2OS cell line with two chromosomally integrated host cell reactivation assays has been previously described (24). Briefly, a U2OS cell line was constructed that contains two chromosomally integrated host cell reactivation assays, one to measure HR and the other to measure mutagenic NHEJ. The functionality of both vectors relies on expression of an I-*Sce*I endonuclease. The I-*Sce*I endonuclease gene is fused to the ligand binding domain of rat glucocorticoid receptor (GR) on the C-terminus, so that the absence of the GR ligand Triamcinolone excludes the I-*Sce*I endonuclease from the nuclease and prevents I-*Sce*I cleavage. To further limit activation of the I-*Sce*I endonuclease, the fusion protein contains a destabilizing domain on the N-terminus. Administration of Shield1 blocks the destabilizing effect. The HR site maintains a Green Fluorescence Reporter (GFP) gene that is interrupted with an I-SceI cleavage site. Upon I-SceI cleavage, a functional GFP gene will be expressed if the cell uses a fragmented and inactivated GFP gene (*iGFP*) on the same or sister chromatid as a template for HR. The NHEJ site maintains a Red Fluorescence Reporter (RFP) gene that is basally repressed by TetR with an I-*Sce*I cleavage site at the TetR locus. If the TetR open reading frame is disrupted, due to I-*Sce*I cleavage and free-end joining during NHEJ, then there is derepression of RFP. Further experimental details are provided in Supplemental Methods.

### Olaparib clonogenic assay

Cells were transfected with miR-24-3p mimic or mimic control. After 24 hours, cells were reseeded and treated with 0 µM, 5 µM, and 10 µM concentrations of Olaparib (Selleckchem) dissolved in DMSO (Sigma). Cells were incubated for 5 days and colonies were stained with a mixture of 6.0% glutaraldehyde (Sigma) and 0.5% crystal violet (Electron Microscopy Sciences). Methanol was added to solubilize the dye. Absorbance at 540 nm was measured using Cytation 3 Cell Imaging Multi-Mode Reader (BioTek).

### Advanced Quantitative analysis (AQUA) for BRCA1 Immunofluorescence

This technique has been previously described (26). Slides were deparaffinized with xylene and rehydrated with ethanol. Antigen retrieval was performed using citrate buffer (pH 6). Non-specific antigens were blocked with 30-minute incubation in 3 % bovine serum albumin in 0.1 mol/L of Tris-buffered saline for 30 min at room temperature. Slides were incubated with primary antibodies for pan-cytokeratin antibody (1:100, Dako) and BRCA1 (antibody validation described previously) at 4°C overnight (1:1000, Abcam) (45), followed by incubation with Alexa 546-conjugated goat anti-rabbit secondary antibody (Life Technologies) diluted 1:100 in mouse EnVision reagent (Dako) for 1 hour. Signal was amplified with Cy5-Tyramide (Perkin Elmer) for 10 minutes and then slides were mounted with ProlongGold + DAPI (Life Technologies). Fluorescent images of DAPI, Alexa 488-cytokeratin and Cy5-BRCA1 were obtained with an Olympus AX-51 epifluorescent microscope. Between 18 and 67 fields of view were quantified for each specimen. Integration of the binarized DAPI and cytokeratin signal generates a compartment image mask. AQUA scores were generated by dividing the sum of the BRCA1 pixel intensities, by the area of the target compartment (DAPI/Cytokeratin image) Analyses was carried out using the AQUAnalysis software (Genoptix).

### Mice and CS exposure

Animal studies were conducted in accordance with the NIH Guide for the Care and Use of Laboratory Animals. AKR/J mice obtained from Jackson Laboratories and bred in our facility. Both female and male; 10-12 weeks old) were used for cigarette smoke (CS) exposure. Littermates were randomized to receiving CS or no CS. Mice were exposed in a Teague TE-10 smoking machine (Teague Enterprises) to CS from 3R4F research cigarettes (University of Kentucky). The number of cigarettes was adjusted between 5-10 per cycle (9 min/cycle) to meet a concentration of 100 mg/m^3^. Cigarette concentration was checked every week. Each smoldering cigarette was puffed for 2 seconds at a flow rate of 1.05 L/min to provide a standard puff of 35 cm^3^. Mice received 6 hours exposure per day, 5 days/week for 5 months, and were sacrificed the day after last CS exposure. Sample collection and morphometric assessments of lungs have been previously described (46). Further details and methods for cleaved caspase 3 immunohistochemistry are provided in Supplemental Methods.

### miR-24-3p locked nucleic acid (LNA) inhibitor and scramble control

Custom miR-24-3p LNA-inhibitor and scramble LNA-control with full phosphorothioate (PS) backbones were designed (Exiqon). Mice were anesthetized with isofluorane and 5 mg/kg of either miR24-3p LNA-inhibitor or LNA-control in 50μl was delivered via intranasal administration. Mice received 1 dose and then were either randomized to CS exposure or non-CS exposure groups. All mice received a subsequent dose after 10 weeks of exposure.

### Statistical Analyses

We used a two-tailed Student’s t-test for two group comparisons of normally distributed continuous data and a Mann-Whitney U test for two group comparisons of non-normally distributed continuous data. A one-way ANOVA was used for multiple group comparisons of normally distributed continuous data and a Kruskal-Wallis test was used for multiple group comparisons of non-normally distributed continuous data. Corrections for multiple comparisons were made using the two-stage step up method of Benjamini, Krieger and Yekutieli. Normality (Gaussian distribution) of the datasets was tested with D’Agostino-Pearson test or Shapiro Wilk normality test. Associations between continuous variables were assessed using a Pearson correlation coefficient for normally distributed data and a Spearman correlation coefficient for non-normally distributed data. For non-array-based studies, a P value of <0.05 was considered significant. For apoptosis assays involving CSE, we accounted for batch effect by normalizing the difference between experimental groups around the mean of all experimental groups. For array-based studies, a false discovery rate (FDR) < 0.05 was considered significant and BRBarray tools was used for analysis (47). For mouse studies, 28 mice (12 males and 16 females) were treated with miR-24-3p LNA-inhibitor and 28 mice (12 males, and 16 females) were treated with LNA-control. All mice were randomly selected amongst littermates amongst 4 breeding cages. Mice were then randomized to receive either CS or no CS, stratifying by gender and treatment group. These sample sizes allowed for 80% power to detect a difference of 17.5% between the two groups at a statistical significance level of 0.05. Actual sample size differed due to mortality or automated detection of erroneous Flexivent measurements. Final sample size was LNA-control + CS (5 males, 6 females) LNA-miR-24-3p inhibitor + CS (6 males, 5 females), LNA-control + No CS (6 males, 7 females), and LNA-miR-24-3p inhibitor + No CS (2 males, 6 females).

### Study approval

All studies on human samples were approved by the Human Investigation Committee (HIC) at Yale University (1409014689) (2000020056). All human tissue samples obtained directly from the LTRC or through the LGRC were collected after ethical review for the protection of human subjects and were approved by the NHLBI-sponsored LTRC. The COSMIC study was approved by the Stockholm regional ethical board number (2006/959-31/1). Written informed consent was received from all participants prior to sample procurement and study inclusion. All animal studies were approved by the Institutional Animal Care and Use Committee (IACUC), Yale University (07867).

## AUTHOR CONTRIBUTIONS

MS generated the main conceptual ideas. JN, JS, PJL, and MS designed research studies. JN, FW, EF, VN, and SK conducted experiments. VN conducted experiments and provided oversight for AQUA of BRCA1. RB provided reagents and oversight for DNA reporter cell line experiments. CMS and AW supervised the COSMIC study. NK supervised data from LGRC. JN, EF, WR, SK, CL, XY and JG analyzed data. MS wrote the manuscript with critical appraisal from NK, PJL, JS, CJB, and AW. All authors contributed to the final version of the manuscript. The order of appearance of co-first authors was determined by seniority and relative contribution to the manuscript.

## ACKNOWLEDGEMENTS

This work was supported by NIH grants K08HL135402, R01HL138396, R01HL127349, U01HL145567, U01HL122626, U54HG008540. and funding from the Swedish Heart Lung-Foundation, Swedish Research Council, Flight Attendant Medical Research Institute, and Claude D. Pepper Older Americans Independence Center, Yale School of Medicine.

## Notes

### Competing Interest Statement

EF is currently employed by Rubius Therapeutics; VN is currently employed by Akoya Bioscience; RB is cofounder and consultant for Cybrexa Therapeutics; MS and PJL are co-inventors on a pending patent describing the therapeutic utility of MIF020 in lung disease; NK reports personal fees from Biogen Idec, Boehringer Ingelheim, Third Rock, Miragen, Pliant, Samumed, NuMedii, Indaloo, Theravance, LifeMax, Optikira, Three Lake Partners and has filed patents related to the use of thyroid hormone as an antifibrotic agent and novel biomarkers in pulmonary fibrosis.

