## Supplemental Figures and Tables for "Decreased miR-24-3p potentiates DNA damage responses and increases susceptibility to COPD"

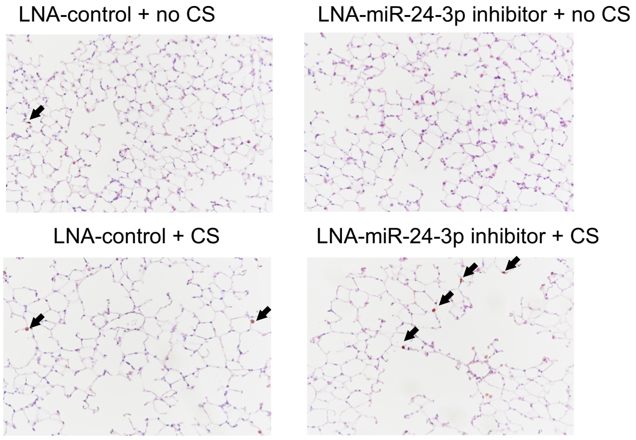


**Supplemental Figure 1** Representative images (20x) of peripheral lung tissue following immunohistochemistry for cleaved caspase 3 in mice treated with locked nucleic acid (LNA) imiR-24-3p inhibitor or LNA-control ± exposure to cigarette smoke (CS). Acquired with 20x objective. Arrows demonstrate cleaved caspase 3 positive cells.

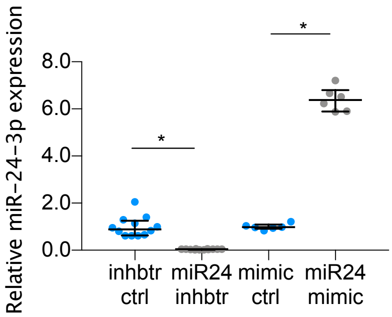


**Supplemental Figure 2** Relative miR-24-3p expression (ΔΔCt miR-24-3p/RNU48) as assessed by RT-PCR following treatment with miR-24-3p mimic vs. mimic control (n=6/group) or miR-24-3p inhibitor vs. inhibitor control (n=12/group) Error bars represent median ± interquartile range. *P<0.05 Mann-Whitney.

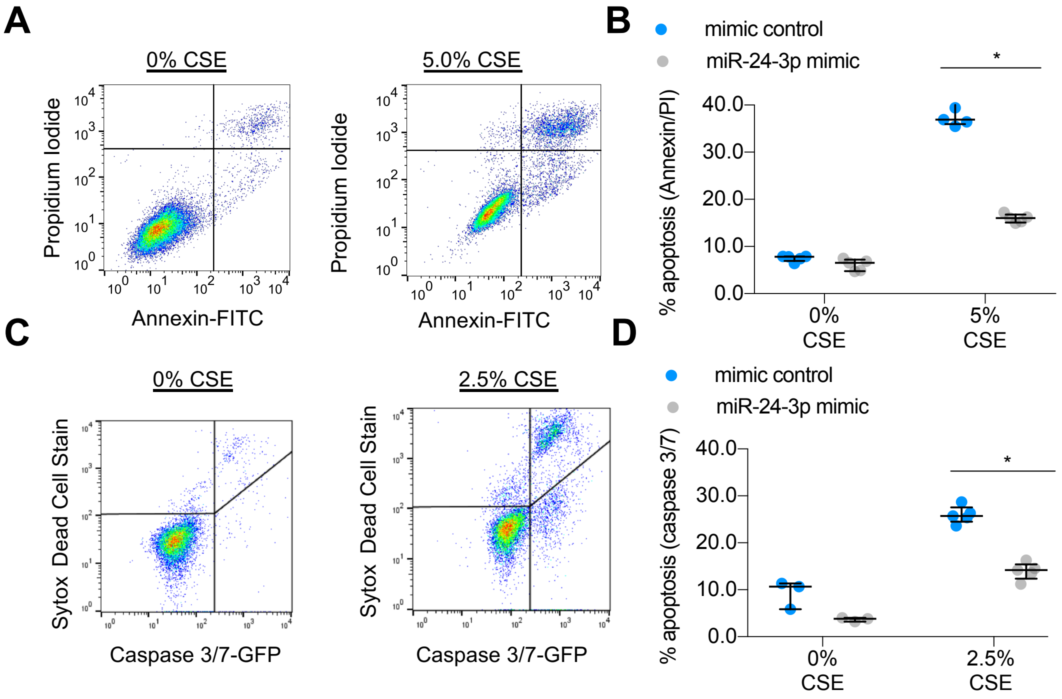


**Supplemental Figure 3 A)** Flow cytometric detection of Propidium Iodide (y-axis) and Annexin V (x-axis) in cells exposed to 0% CSE (*left*) or 5% cigarette smoke extract (CSE) *(right)* **B**) Percent apoptotic cells determined by Annexin/PI staining in miR-24-3p mimic vs. mimic control treated BEAS2B cells exposed to 0%, and 5% cigarette smoke extract (CSE). n=5/group. **C** Flow cytometric detection of Sytox AAdvanced(y-axis) and caspase 3/7 (x-axis) in cells exposed to 0% CSE (*left*) or 2.5% cigarette smoke extract (CSE) (*Right*). **D)** Percent apoptotic cells determined by caspase 3/7 and Sytox in miR-24-3p mimic and mimic control treated BEAS2B cells exposed to 0% or 2.5% CSE. n=4-5/group. Error bars represent median ± interquartile range. *P<0.05 Kruskal-Wallis correcting for multiple comparisons using 2-stage linear step-up procedure of Benjamini, Krieger, and Yekutieli.

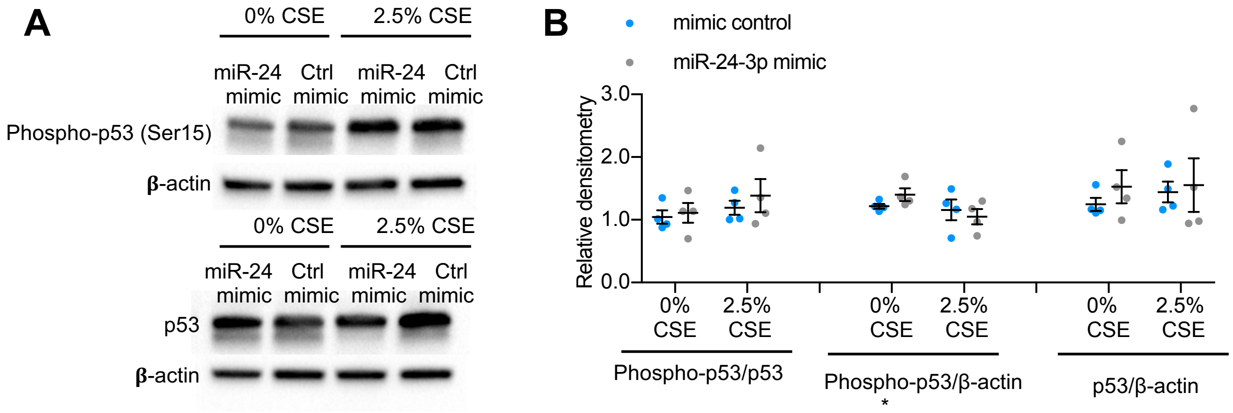


**Supplemental Figure 4 A)** Sample immunoblotting for phosphorylated p53 (Serine 15), p53, and β-actin performed on lysates of BEAS2B cells treated with miR-24-3p mimic, mimic control, miR-24-3p inhibitor, or inhibitor control ± cigarette smoke extract (CSE). **B)** Relative densitometry of phosphorylated p53 (Serine 15), p53, and β-actin. n=4/group. Error bars represent median ± interquartile range.

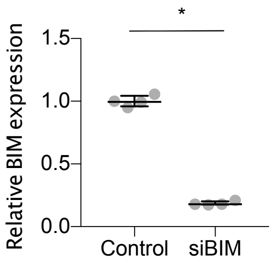


**Supplemental Figure 5** Relative BIM expression in BEAS2B cells following treatment with scrambled control or silencing RNA targeted against BIM (siBIM). n=4/group. Error bars represent median ± interquartile range. *P<0.05 using Mann-Whitney.

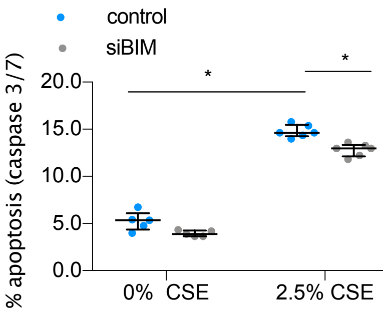


**Supplemental Figure 6** Percent apoptotic cells determined by flow cytometry for caspase 3/7 and Sytox in BEAS2B treated with scrambled control or silencing RNA targeted against BIM (siBIM) and exposed to 0% or 2.5% CSE. n=5-6/group. Error bars represent median ± interquartile range. *P<0.05, Kruskal-Wallis correcting for multiple comparisons using 2-stage linear step-up procedure of Benjamini, Krieger, and Yekutieli.

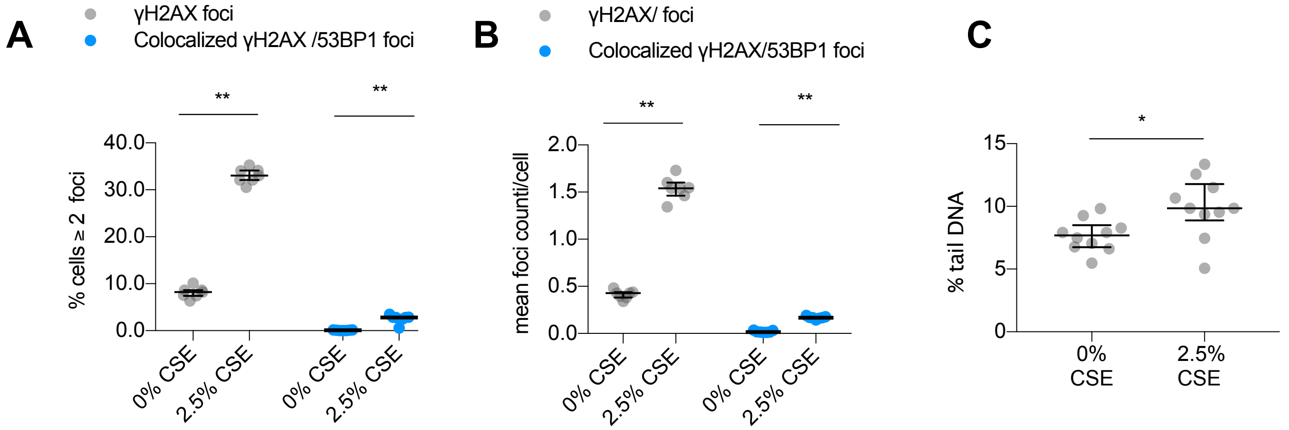


**Supplemental Figure 7 A**) Number of **γ**H2AX foci (grey) or colocalized **γ**H2AX and 53BP1 foci (blue) in BEAS2B cells exposed to 2.5% CSE for 24 hours. n=7/group. **B)** Percent of BEAS2B cells with ≧ 2 **γ**H2AX foci (grey) or colocalized **γ**H2AX and 53BP1 foci (blue) over time. n=7/group. **C)** Results of Comet assay in BEAS2B cells exposed to 0% or 2.5% cigarette smoke extract (CSE) for 24 hours. Percent tail DNA reflects increased DNA damage. n=10/group. Error bars represent median ± interquartile range. **P<0.001, *P<0.05 Mann-Whitney.

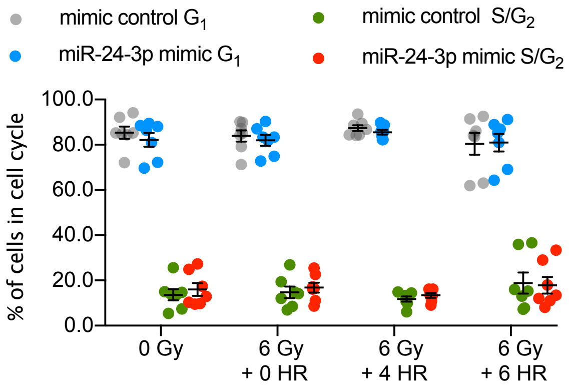


**Supplemental Figure 8** Cell cycle phase in primary human airway epithelial cells as estimated by DAPI staining. n=7/group. Error bars represent median ± interquartile range.

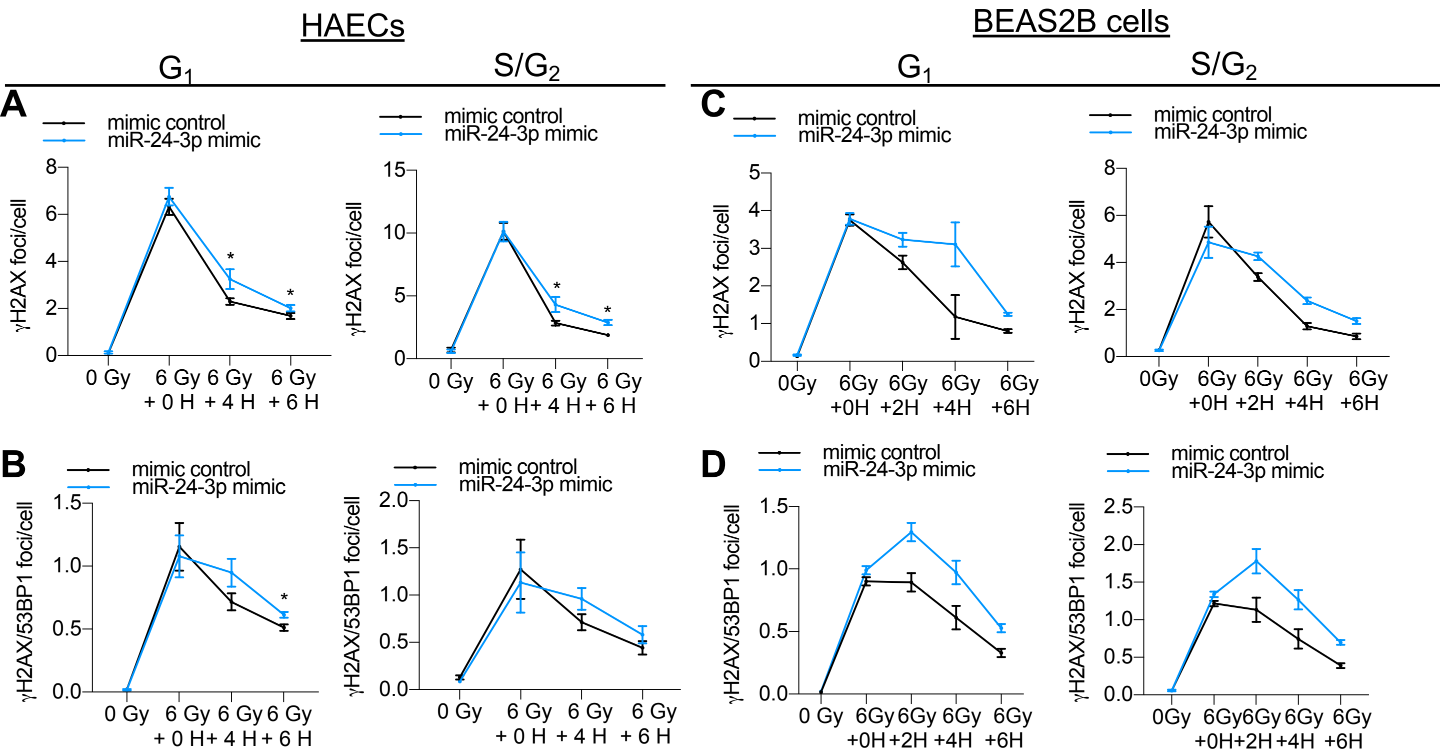


**Supplemental Figure 9 A-D)** Primary human airway epithelial cells (HAECs) or BEAS2B (exposed to 0 Grey (Gy) ionizing radiation or 6 Gy with 0-6 hours (H) of recovery to assess kinetics of DNA repair **A**) Number of **γ**H2AX foci in HEBCs over time in $G_1$ or $S/G_2$ phase of cell cycle. **B)** Number of colocalized **γ**H2AX and 53BP1 foci in HAECs over time in $G_1$ or $S/G_2$ of cell cycle **C**) Number of **γ**H2AX foci in BEAS2B over time in $G_1$ or $S/G_2$ phase of cell cycle **.D)** number of colocalized **γ**H2AX and 53BP1 foci in BEAS2B over time in $G_1$ or $S/G_2$ of cell cycle. Error bars represent mean ± SEM (**A, B**) or median ± interquartile range (**C, D**). *P<0.05 ordinary one-way ANOVA (**A,B**) or Kruskal-Wallis (**C,D**) correcting for multiple comparisons using 2-stage linear step-up procedure of Benjamini, Krieger, and Yekutieli.

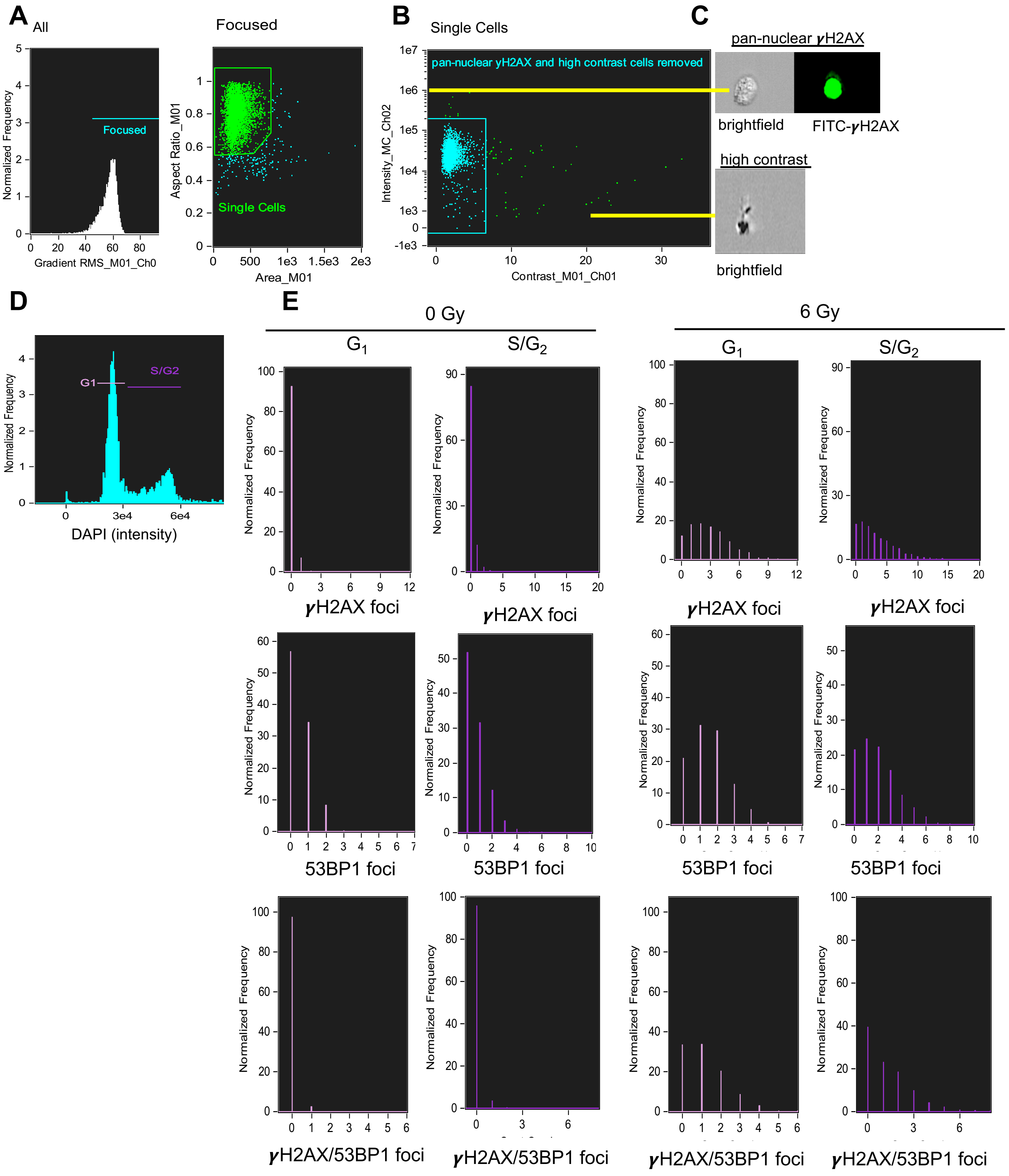


**Supplemental Figure 10 A)** We used the Gradient RMS feature in the brightfield channel, which indicate sharpness of an image, to identify cells in focus. **B**) We used the area and aspect ratio features in the brightfield channel to identify single cells and remove doublets and debris. **C**) We use the intensity feature in channel 2 to identify pan nuclear γH2AX staining and the contrast feature in the brightfield channel to identify high contrast cells. Both features are suggestive of dead/dying cells. **D**) Cell cycle was estimated from DNA staining with DAPI. **E**) Automated counting of foci of γH2AX , 53BP1, and combined γH2AX/53BP1 was conducted using Imagine software. A distribution of these foci in $G_1$ and S/$G_2$ phase following 0 Grey (Gy) or 6 Gy of radiation are shown.

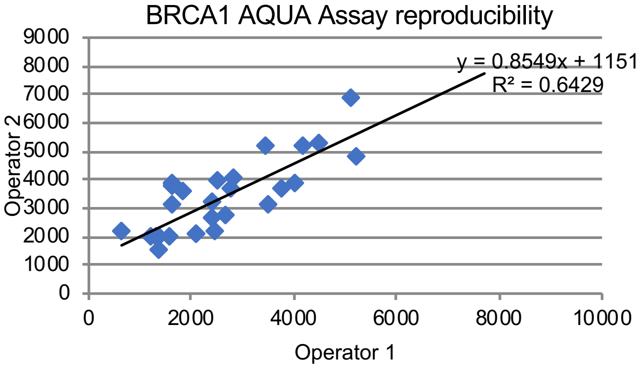


**Supplemental Figure 11** To confirm reliability of the BRCA1 AQUA technique, the assay was performed by two independent operators on two separate days. n= 23.

**Supplemental Table 1** Spearman correlations of microRNAs with FEV$_1$ percent predicted (*left*) and percent radiographic emphysema (*right*) with a minimum P-value > 0.05. FDR = False Discovery Rate.

| Spearman Correlation with FEV1 | | | |
|---|---|---|---|
| Correlation coefficient | Parametric p-value | FDR | UniqueID |
| 0.424 | < 1e-07 | < 1e-07 | hsa-miR-181d-5p |
| 0.378 | 5.00E-07 | 9.65E-05 | hsa-miR-551b-3p |
| 0.343 | 5.40E-06 | 0.000695 | hsa-miR-24-3p |
| 0.336 | 8.30E-06 | 0.000801 | hsa-miR-30d-5p |
| 0.326 | 1.64E-05 | 0.00127 | hsa-miR-30b-5p |
| -0.316 | 2.98E-05 | 0.00192 | hsa-miR-21-5p |
| 0.309 | 4.44E-05 | 0.00245 | hsa-miR-203-3p |
| 0.302 | 6.85E-05 | 0.00331 | hsa-miR-30a-5p |
| -0.297 | 9.19E-05 | 0.00394 | hsa-miR-148a-3p |
| 0.29 | 0.0001345 | 0.00501 | hsa-miR-181c-5p |
| -0.289 | 0.0001428 | 0.00501 | hsa-miR-4306 |
| -0.284 | 0.0001839 | 0.00592 | hsa-miR-21-3p |
| 0.274 | 0.0003161 | 0.00939 | hsa-miR-181b-5p |
| 0.271 | 0.0003657 | 0.0101 | hsa-miR-151-3p |
| 0.268 | 0.0004388 | 0.0113 | hsa-miR-30c-2-3p |
| 0.264 | 0.00052 | 0.0125 | hsa-miR-30a-3p |
| 0.257 | 0.0007277 | 0.0157 | hsa-miR-27a-3p |
| -0.257 | 0.0007311 | 0.0157 | hsa-miR-199a-5p |
| 0.255 | 0.00081 | 0.0165 | hsa-miR-338-3p |
| -0.252 | 0.0009592 | 0.0185 | hsa-miR-142-3p |
| 0.246 | 0.0012341 | 0.0227 | hsa-miR-455-5p |
| 0.241 | 0.0015924 | 0.0279 | hsa-miR-181a-5p |
| 0.229 | 0.0026854 | 0.0451 | hsa-miR-210-3p |
| 0.224 | 0.0033658 | 0.0541 | hsa-miR-598-3p |
| 0.221 | 0.003784 | 0.0584 | hsa-miR-4317 |
| 0.219 | 0.0041403 | 0.06 | hsa-miR-126-3p |
| -0.219 | 0.0041947 | 0.06 | hsa-miR-223-3p |
| -0.216 | 0.0048161 | 0.0664 | hsa-miR-144-3p |
| 0.214 | 0.0050626 | 0.0674 | hsa-miR-498 |
| -0.21 | 0.0060417 | 0.0762 | hsa-miR-185-5p |
| 0.21 | 0.0061181 | 0.0762 | hsa-miR-455-3p |
| 0.209 | 0.0063324 | 0.0764 | hsa-miR-3065-5p |
| -0.207 | 0.0068105 | 0.0774 | hsa-miR-582-5p |
| -0.207 | 0.0069353 | 0.0774 | hsa-miR-142-5p |
| 0.206 | 0.007109 | 0.0774 | hsa-miR-331-3p |
| -0.206 | 0.0072176 | 0.0774 | hsa-miR-425-3p |
| 0.205 | 0.0075385 | 0.0786 | hur_6 |
| -0.204 | 0.0078614 | 0.0799 | hsa-miR-214-3p |
| 0.2 | 0.008842 | 0.0843 | hsa-miR-125a-5p |
| 0.2 | 0.0091506 | 0.0843 | hsa-miR-30c-5p |
| -0.2 | 0.0091566 | 0.0843 | hsa-miR-1246 |
| -0.2 | 0.0091705 | 0.0843 | hsa-miR-199a-3p |
| 0.197 | 0.0100353 | 0.0893 | hsa-miR-30b-3p |
| -0.197 | 0.0101837 | 0.0893 | hsa-miR-144-3p |
| -0.196 | 0.0104942 | 0.09 | hsa-miR-100-5p |
| 0.193 | 0.0117325 | 0.0985 | hsa-miR-218-5p |
| 0.192 | 0.0122758 | 0.099 | hsa-miR-130a-3p |
| 0.192 | 0.0123072 | 0.099 | hsa-miR-151a-5p |
| 0.191 | 0.0125717 | 0.099 | hsa-miR-197-3p |
| 0.186 | 0.0153498 | 0.119 | hsa-miR-326 |
| -0.185 | 0.016064 | 0.12 | hsa-miR-19a-3p |
| -0.184 | 0.0161451 | 0.12 | hsa-miR-376a-3p |
| -0.183 | 0.0168351 | 0.121 | hsa-miR-495-3p |
| 0.183 | 0.0169747 | 0.121 | hsv1-miR-H8-5p |
| 0.182 | 0.0178023 | 0.123 | hsa-miR-4270 |
| -0.182 | 0.0179098 | 0.123 | hsa-miR-584-5p |
| 0.181 | 0.0181485 | 0.123 | hsa-miR-187-5p |
| 0.18 | 0.0186406 | 0.124 | hsa-miR-1181 |
| 0.18 | 0.0190584 | 0.125 | hsa-miR-224-5p |
| -0.179 | 0.0195736 | 0.126 | hsa-miR-146b-5p |
| 0.178 | 0.0199528 | 0.126 | hsa-miR-23a-3p |
| 0.176 | 0.0219107 | 0.136 | hsa-miR-4322 |
| -0.175 | 0.0229059 | 0.14 | hsa-miR-3651 |
| 0.174 | 0.0232032 | 0.14 | hsa-miR-99b-5p |
| -0.173 | 0.0241515 | 0.141 | hsa-miR-136-3p |
| -0.173 | 0.0242629 | 0.141 | hsa-miR-125b-5p |
| 0.172 | 0.0248808 | 0.141 | hsv2-miR-H24 |
| 0.172 | 0.0250556 | 0.141 | hsa-miR-34c-5p |
| -0.172 | 0.0252105 | 0.141 | hsa-miR-16-5p |
| 0.171 | 0.0257163 | 0.142 | hsa-miR-34b-5p |
| 0.17 | 0.0262983 | 0.143 | hsa-miR-574-3p |
| -0.17 | 0.0268006 | 0.144 | hsa-miR-34a-5p |
| -0.169 | 0.0280249 | 0.148 | hsa-miR-4284 |
| -0.167 | 0.0300203 | 0.155 | hsa-miR-140-5p |
| 0.167 | 0.0300276 | 0.155 | hsa-miR-1249-3p |
| 0.165 | 0.0316558 | 0.161 | hsa-miR-1226-5p |
| 0.164 | 0.0324755 | 0.163 | hsa-miR-30d-3p |
| 0.161 | 0.035615 | 0.175 | hsa-miR-296-5p |
| 0.161 | 0.0359346 | 0.175 | hsa-miR-452-5p |
| -0.161 | 0.036357 | 0.175 | hsa-miR-199b-5p |
| 0.16 | 0.0376937 | 0.18 | hsa-miR-26a-5p |
| 0.157 | 0.0408673 | 0.19 | hsa-miR-141-3p |
| -0.157 | 0.0409179 | 0.19 | hsa-miR-376c-3p |
| 0.156 | 0.0428897 | 0.197 | hsa-miR-221-3p |
| 0.154 | 0.0443135 | 0.199 | hsv2-miR-H22 |
| 0.154 | 0.0447422 | 0.199 | hsa-miR-33a-5p |
| -0.154 | 0.0448448 | 0.199 | hsa-miR-451a |
| -0.152 | 0.047776 | 0.21 | hsa-miR-15a-5p |
| 0.151 | 0.0491452 | 0.213 | hsa-miR-3188 |

| Spearman Correlation with percent emphysema | | | |
|---|---|---|---|
| Correlation coefficient | Parametric p-value | FDR | UniqueID |
| -0.353 | 0.000131 | 0.0231 | hsa-miR-24-3p |
| -0.347 | 0.0001684 | 0.0231 | hsa-miR-30a-5p |
| -0.346 | 0.0001797 | 0.0231 | hsa-miR-181d-5p |
| 0.301 | 0.0011744 | 0.0967 | hsa-miR-199a-5p |
| -0.296 | 0.001453 | 0.0967 | hsa-miR-551b-3p |
| -0.295 | 0.0015027 | 0.0967 | hsa-miR-338-3p |
| 0.285 | 0.0021706 | 0.101 | has-miR-144-3p |
| 0.283 | 0.0023347 | 0.101 | hsa-miR-4306 |
| -0.283 | 0.0023656 | 0.101 | hsa-miR-181a-5p |
| -0.272 | 0.0035378 | 0.123 | hsa-miR-331-3p |
| 0.268 | 0.0039881 | 0.123 | hsa-miR-451a |
| -0.266 | 0.004268 | 0.123 | hsa-miR-203-3p |
| -0.266 | 0.0043811 | 0.123 | hsa-miR-181b-5p |
| -0.265 | 0.0045259 | 0.123 | hur_6 |
| -0.263 | 0.0047964 | 0.123 | hsa-miR-30b-5p |
| 0.26 | 0.0053106 | 0.128 | hsa-miR-199a-3p |
| -0.256 | 0.0060748 | 0.138 | hsa-miR-30d-5p |
| -0.251 | 0.0072342 | 0.155 | hsa-miR-200c-3p |
| 0.248 | 0.0078927 | 0.16 | hsa-miR-144-3p |
| -0.244 | 0.0089303 | 0.172 | hsa-miR-598-3p |
| -0.241 | 0.0100427 | 0.184 | hsa-miR-197-3p |
| 0.238 | 0.0108938 | 0.184 | hsa-miR-486-5p |
| -0.238 | 0.0109631 | 0.184 | hsa-miR-151a-5p |
| -0.236 | 0.0117602 | 0.184 | hsa-miR-181a-3p |
| 0.235 | 0.0118969 | 0.184 | hsa-miR-223-3p |
| 0.231 | 0.0135586 | 0.201 | hsa-miR-21-3p |
| -0.23 | 0.0140969 | 0.202 | hsa-miR-33a-5p |
| -0.228 | 0.014834 | 0.204 | hsa-miR-30a-3p |
| -0.226 | 0.0155854 | 0.207 | hsa-miR-126-3p |
| -0.223 | 0.0173123 | 0.223 | hsa-miR-30c-5p |
| 0.217 | 0.0203321 | 0.249 | hsa-miR-198 |
| 0.217 | 0.0207598 | 0.249 | hsa-miR-483-5p |
| -0.216 | 0.021309 | 0.249 | hsa-miR-326 |
| -0.213 | 0.0229003 | 0.26 | hsa-miR-30e-3p |
| -0.212 | 0.0240042 | 0.261 | hsa-miR-181c-5p |
| -0.211 | 0.0243525 | 0.261 | hsa-miR-151-3p |
| -0.207 | 0.0271023 | 0.283 | hsa-miR-30c-2-3p |
| -0.206 | 0.0282946 | 0.287 | dmr_3 |
| -0.204 | 0.0292634 | 0.29 | hsa-miR-1271-5p |
| 0.2 | 0.032661 | 0.315 | hsa-miR-214-3p |
| -0.199 | 0.0337918 | 0.318 | hsa-miR-30b-3p |
| -0.198 | 0.0353433 | 0.325 | hsa-miR-3065-5p |
| 0.196 | 0.0366227 | 0.326 | hsa-miR-320c |
| -0.195 | 0.0376419 | 0.326 | dmr_6 |
| -0.195 | 0.0380465 | 0.326 | hsa-miR-27a |
| 0.194 | 0.0392238 | 0.329 | hsa-miR-185-5p |
| -0.19 | 0.043302 | 0.356 | hsa-miR-542-5p |
| -0.189 | 0.0443359 | 0.357 | hsa-miR-1288-3p |
| 0.188 | 0.0453903 | 0.358 | hsa-miR-16-5p |
| 0.186 | 0.0476306 | 0.368 | hsa-miR-16-2-3p |

**Demographic characteristics of study patients - LGRC cohort (Discovery and Validation cohorts)**

| | Discovery cohort | | | Validation cohort | | |
|---|---|---|---|---|---|---|
| | **No COPD** | **GOLD I, II** | **GOLD III, IV** | **No COPD** | **GOLD I, II** | **GOLD III, IV** |
| Subjects | 28 | 36 | 20 | 50 | 14 | 24 |
| Female sex (%) | 15 (54%) | 17 (47%) | 11 (55%) | 30 (59%) | 7 (50%) | 14 (58%) |
| Age | 64 (56 - 73) | 70 (65 - 74) | 64 (57 - 70) | 66 (60 - 72) | 64 (60 - 73) | 62 (51 - 67) |
| Race or ethnic group | 23 W,1 H,1 AA,1A,1O | 35 W, 1 AA | 19 W, 1 O | 50 W, 1 AA | 13 W, 1 AA | 23 W, 1 AA |
| $FEV_1$ % predicted | 97 (87 - 105) | 62 (55 - 73)[1] | 25 (22 - 33)[ [1] | 91 (85 - 101) | 59 (50 - 72) | 22 (16 - 28) |
| % emphysema | 0.14(0.09-0.45) [25] | 2.55(0.78-11.20) | 25.50(14.90-37.80)[2] | 0.26(0.14-0.54) [23] | 4.45(0.87-9.00) | 25.90(18.00-32.10) |
| smoking status | 15 F, 9 N [4] | 31 F, 1 N, 4 C | 18 F, 1 N, 1 C | 31 N, 14 F, 1 C [5] | 14 F | 23 F, 1 C |
| Pack-Year | 7 (0 - 26) [6] | 60 (31 - 115) | 50 (42 - 80) | 17 (0 - 47) [5] | 50 (40 - 80) | 40 (21 - 59) [1] |

**Supplemental Table 2** LGRC = Lung Genomics Research Consortium, GOLD = Global initiative for Obstructive Lung Disease, W=White, H=Hispanic, AA=African American, A=Asian, O=other, F=former smoker, N=never smoker, C=current smoker. [] represents missing data. () represents interquartile range.

**Demographic characteristics of study patients – Confirmatory cohort**

| | **GOLD 0** | **GOLD I, II** | **GOLD III, IV** |
|---|---|---|---|
| Subjects | 28 | 35 | 24 |
| Female sex (%) | 11 (39%) | 14 (40%) | 11 (46%) |
| Age | 65 ±13 | 71±9 | 58±12 |
| Race or ethnic group | 28 W | 35 W | 22 W 2 AA |
| $FEV_1$ percent predicted | 96 ± 14 | 69 ± 15 | 26 ± 6 |
| % radiographic emphysema | 0.25 (0.13 - 2.26) [12] | 3.7 (1.00 - 7.30) | 25.80 (9.10 – 36.40) |
| smoking status | 19F, 7N [2] | 32 F, 2N, 1C | 24F |
| Pack-Year | 12 (0 – 53) [2] | 50 (37 – 77) | 42 (20 - 65)[1] |

**Supplemental Table 3** LGRC = Lung Genomics Research Consortium, GOLD = Global initiative for Obstructive Lung Disease W=White, AA=African American, F=former smoker, N=never smoker, C=current smoker.[] represents missing data. () represents interquartile range. ± represents standard deviation.

**Demographic characteristics of study patients – COSMIC Cohort**

| | **Never smokers** | **Smokers without COPD** | **Smokers with COPD (GOLD I)** | **Smokers with COPD (GOLD II)** |
|---|---|---|---|---|
| Subjects | 22 | 10 | 18 | 13 |
| Female sex (%) | 11 (50%) | 3 (30%) | 10 (56%) | 5 (38%) |
| Age | 59 (47 - 62) | 55 (50 - 62) | 61 (54 - 64) | 63 (59 - 65) |
| $FEV_1$ percent predicted | 116 ± 14 | 107 ± 10 | 87 ± 5 | 69 ± 9 |
| Pack-year | - | 35 ± 7 | 36 ± 9 | 42 ± 15 |
| Smoking status | - | 10 C | 12 C, 6 F | 8 C, 5 F |

**Supplemental Table 4** COSMIC =Clinical & Systems Medicine Investigations of Smoking-related Chronic Obstructive Pulmonary Disease) GOLD = Global initiative for Obstructive Lung Disease. F=former smoker, N=never smoker, C=current smoker. [] represents missing data. () represents interquartile range. ± represents standard deviation.

| Demographic characteristics of study patients – BIM western blot | | |
|---|---|---|
| | **GOLD I** | **GOLD III, IV** |
| Subjects | 6 | 9 |
| Female sex (%) | 2 (44.44%) | 4 (44.44%) |
| Age | 71 (68 - 82) | 59 (56 - 65) |
| Race or ethnic group | 5 W, 1 AA | 8 W, 1 AA |
| $FEV_1$ percent predicted | 93 (79 - 100) | 21 (19 - 25) |
| Smoking status | 6 F | 9 F |
| Pack-year | 38 (20 - 69) | 38 (30 - 54) |

**Supplemental Table 5** Demographic characteristics of study patients from which tissue was obtained for immunoblotting for BIM. GOLD = Global initiative for Obstructive Lung Disease. E = ever smoker [] represents missing data. () represents interquartile range.

| Demographic characteristics of study patients - BRCA1 AQUA staining | | |
|---|---|---|
| | **NO COPD** | **COPD** |
| Subjects | 9 | 16 |
| Female sex (%) | 4 (44%) | 4 (25%) |
| Age | 62 ± 14 | 63 ± 8 |
| Race or ethnic group | 9 W | 14 W, 2 AA |
| $FEV_1$ percent predicted | 89 (82 - 94) | 25 (22 - 35) |
| % radiographic emphysema | 0.03 (0 – 0.19) | 30.20 (17.20 – 43.10) |
| Smoking status | 8 F, 1 N | 16 F |
| Pack-Year | 19 (2 - 46) [1] | 43 (32 - 74) |

**Supplemental Table 6** Demographic characteristics of study patients from which tissue was obtained for BRCA1 AQUA staining. W = white, AA = African American. E=ever smoker, N= never smoker. [] represents missing data. () represents interquartile range. ± represents standard deviation

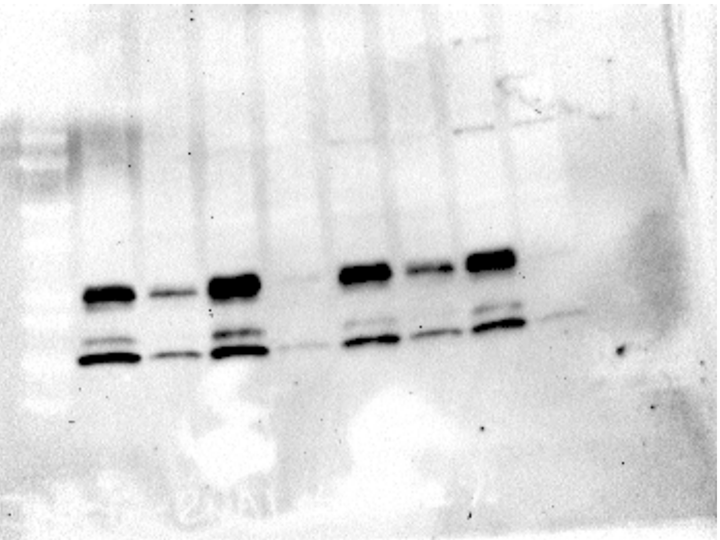

Figure 3E - BIM

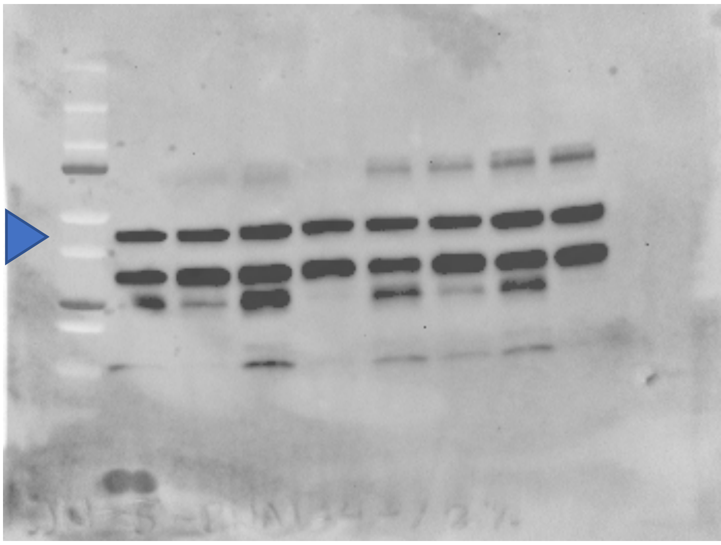

Figure 3E - **β**-actin

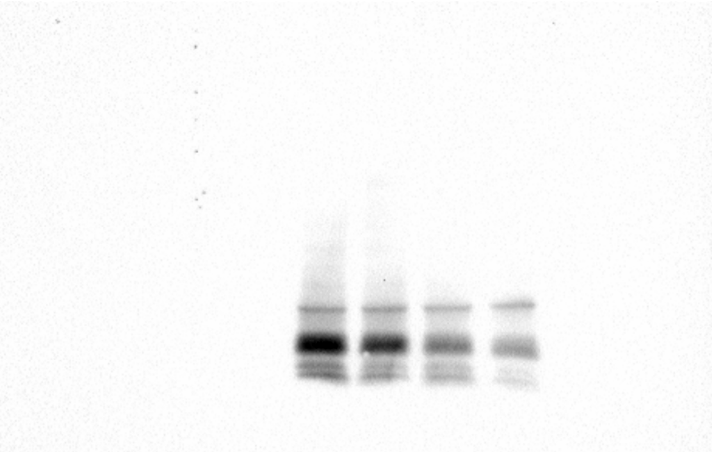

Figure 3E - BIM

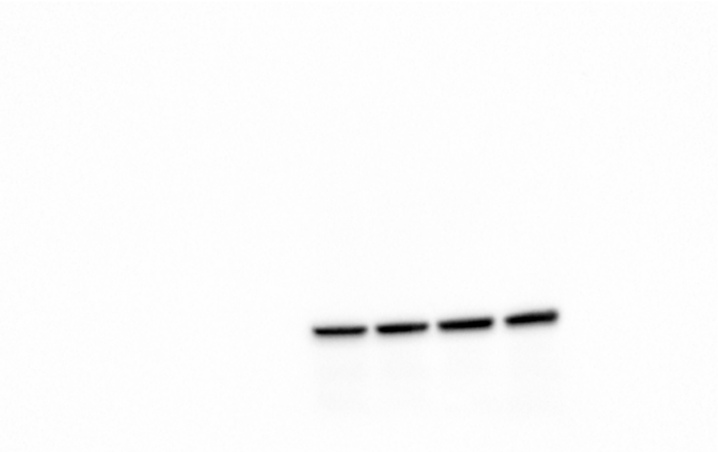

Figure 3E -BIM

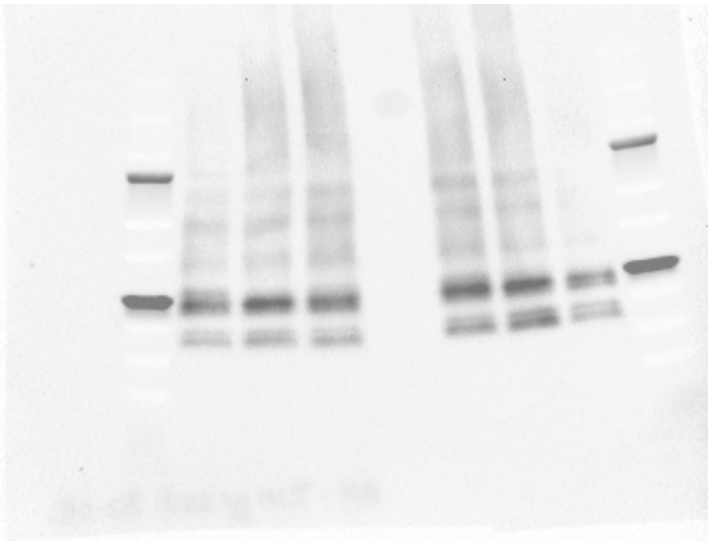

Figure 3H - BIM (RA)

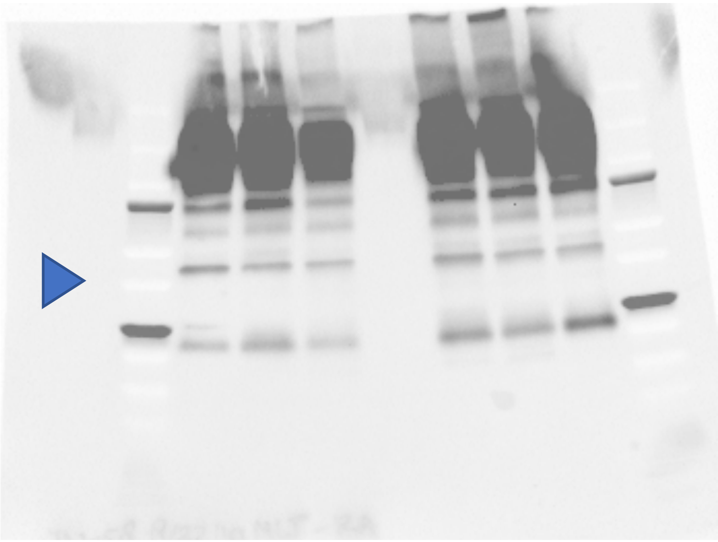

Figure 3H - **β**-actin (RA)

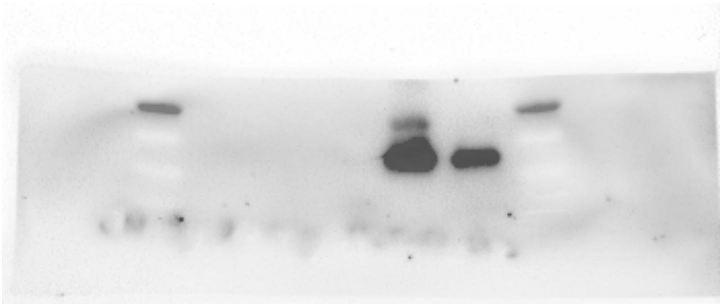

Figure 3H - BIM (CS)

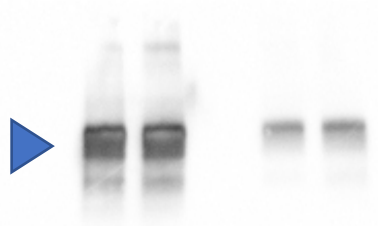

Figure 3H - **β**-actin (CS)

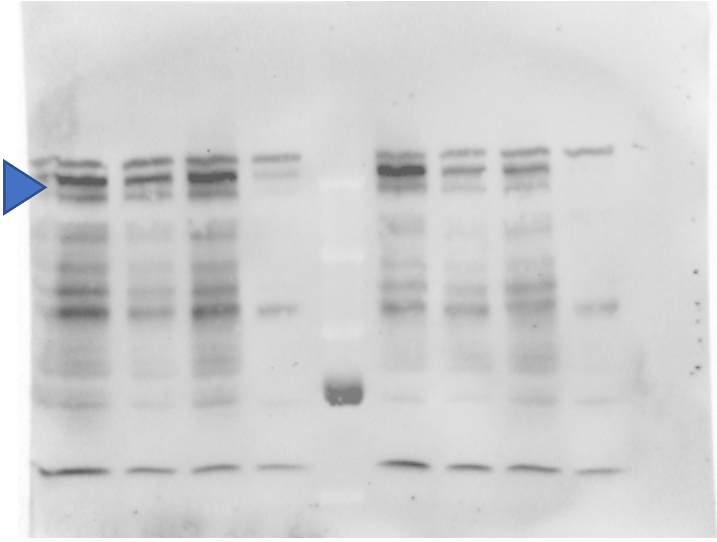

4K - BRCA1

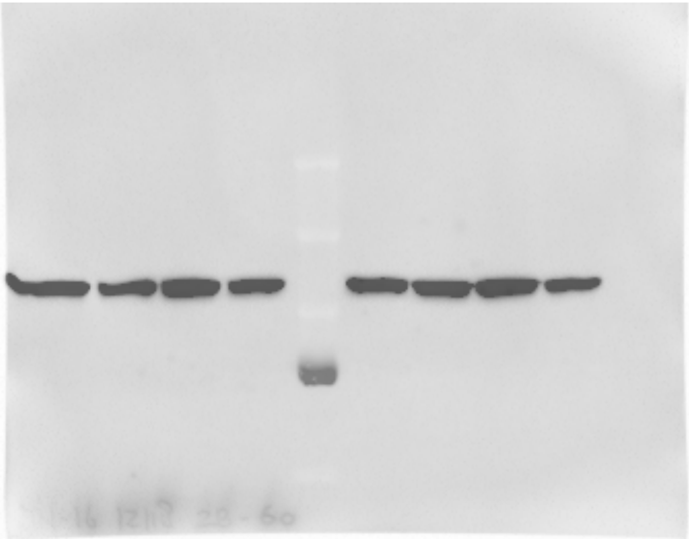

4K - β-actin

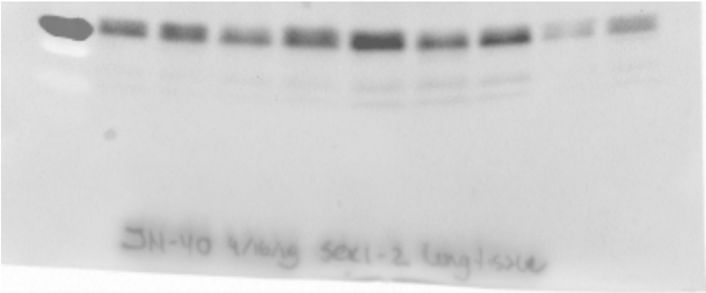

5D - BIM

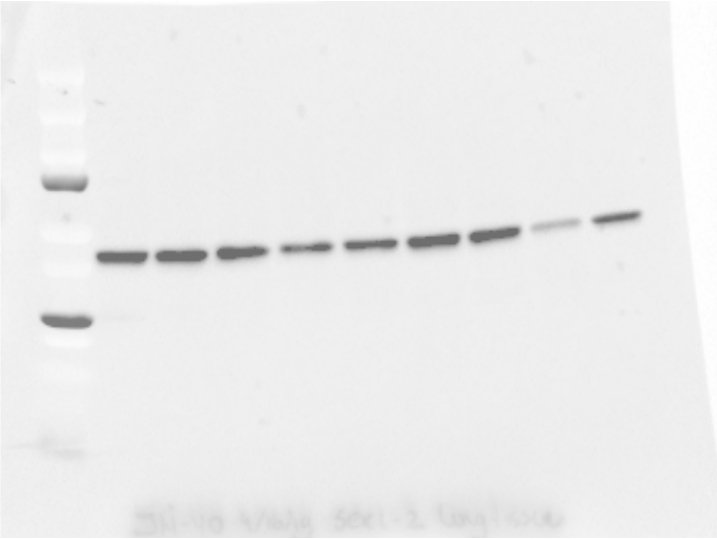

5D - β-actin

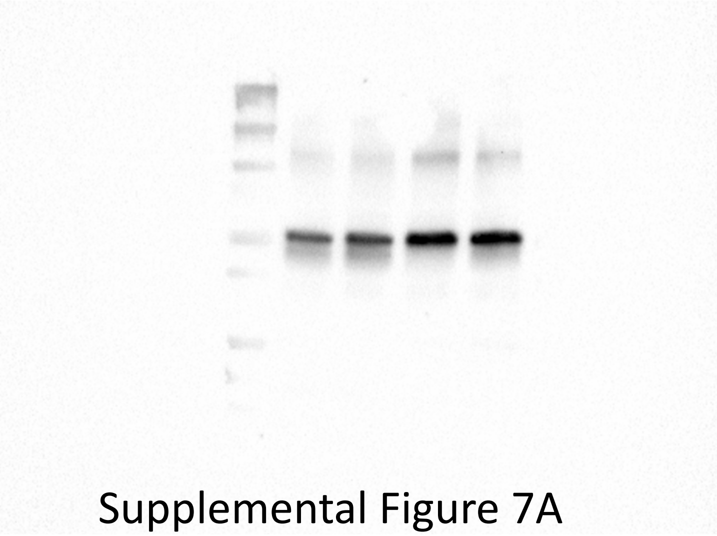

Supplemental Figure 7A
phospho-p53

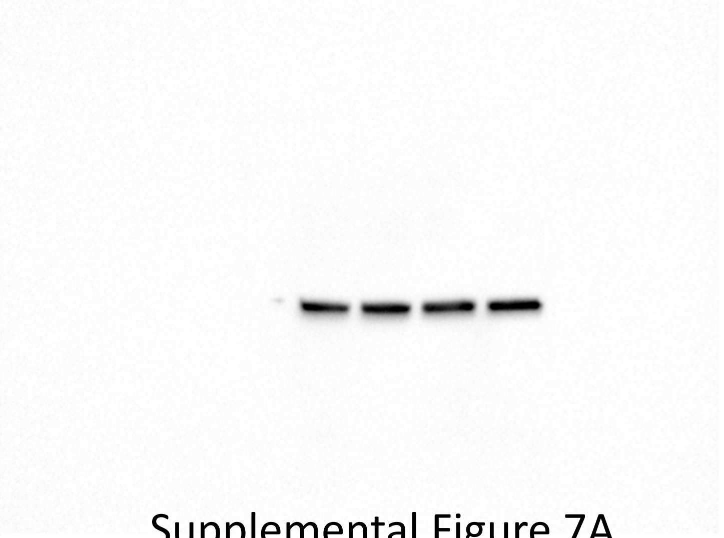

Supplemental Figure 7A
**β**-actin

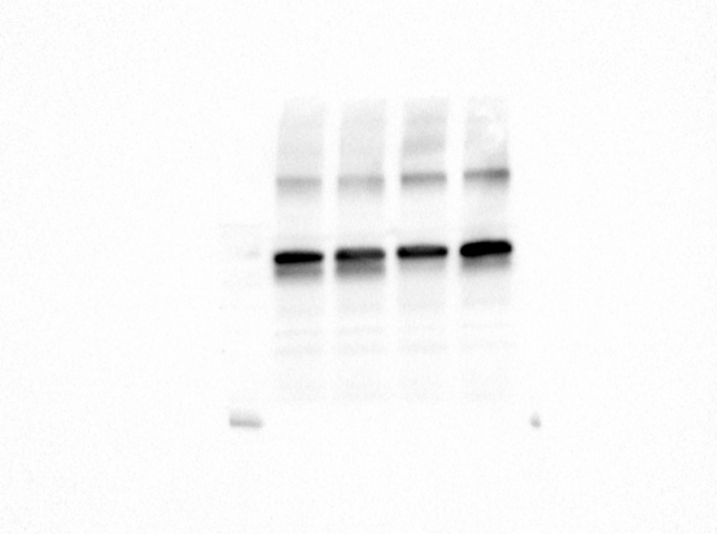

Supplemental Figure 7A
p53

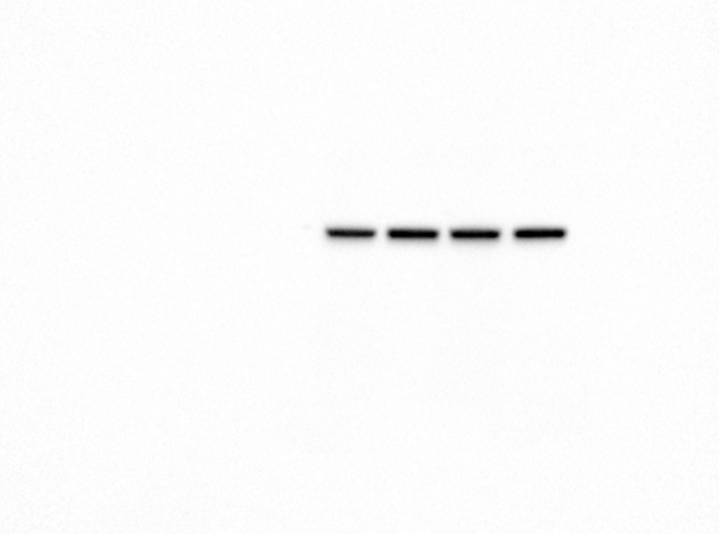

Supplemental Figure 7A
**β**-actin
